# Influence of genetic ancestry on breast stromal cells provides biologic basis for increased incidence of metaplastic breast cancer in women of African descent

**DOI:** 10.1101/2022.07.14.500115

**Authors:** Brijesh Kumar, Katie Batic, Poornima Bhat-Nakshatri, Maggie M Granatir, Rebekah Joann Addison, Megan Szymanski, Lee Ann Baldridge, Constance J. Temm, George Sandusky, Sandra K Althouse, Anna Maria Storniolo, Harikrishna Nakshatri

## Abstract

The biologic basis of genetic ancestry-dependent variability in disease incidence and outcome is just beginning to be explored. We recently reported enrichment of a population of ZEB1-expressing cells located adjacent to the ductal epithelial cells in the normal breast of women of African Ancestry (AA) compared to European Ancestry (EA). By establishing and characterizing cell lines corresponding to these cells and validating *in vitro* findings with tissue microarrays of healthy breast tissue from AA, EA and Latina Ancestry (LA) women, we demonstrate that these cells have the properties of fibroadipogenic/mesenchymal stromal cells that express PROCR and PDGFRα. <u>P</u>ROCR+/<u>Z</u>EB1+/<u>P</u>DGFRα+ cells, hence renamed as PZP cells, are enriched in the normal breast tissues of AA compared to EA or LA women. *In vitro*, PZP cells trans-differentiated into adipocytes or osteocytes. In co-culture conditions, PZP:epithelial cell communication resulted in luminal epithelial cells acquiring basal/stem cell characteristics and increased expression of IL-6 suggesting the impact of this communication on breast epithelial hierarchy and the microenvironment. Consistent with this possibility, the level of phospho-STAT3, which is a downstream target of IL-6, was higher in the normal and cancerous breast tissues of AA compared to EA women. PZP cells transformed with HRas^G12V^ ± SV40-T/t antigens generated metaplastic carcinoma in NSG mice suggesting that these cells could be the cell-of-origin of metaplastic breast cancers. Collectively, these results identify a stromal cell component that could influence the biology of breast cancer in AA women.

## INTRODUCTION

Mortality from triple negative breast cancer (TNBC) is much higher in women of African Ancestry (AA) compared to women of European Ancestry (EA) even when controlled for socioeconomic status ^1^. Even in cases of estrogen receptor positive (ER+)/HER2-/node negative breast cancers with intermediate Oncotype recurrence score ^2^, clinical outcome is worse in AA women compared to EA women ^3^. Furthermore, ductal carcinoma in situ (DCIS), which is typically considered to be of minimal risk of recurrence, tends to be more deleterious to AA women compared to EA women ^4^. With the recent discovery of the presence of an extra 296,485,284 base pairs DNA in populations of African descent affecting 315 distinct protein-coding genes ^5^, the disparity in breast cancer outcome could, therefore, partly be due to genetic ancestry-dependent variability in normal and cancerous breast biology.

To discover genetic ancestry-dependent differences in normal breast biology, we initiated studies using healthy breast tissues of women of different genetic ancestry donated to the Susan G. Komen Tissue Bank (KTB) at the IU Simon Comprehensive Cancer Center (IUSCCC). We had previously demonstrated that cultured cells from breast tissues of AA women are enriched for PROCR^+^/EpCAM^-^ and CD44^+^/C24^-^ cells compared to EA women^6^. In the mouse mammary gland, the PROCR^+^/EpCAM^‒^ cells have been demonstrated to function as multi-potent stem cells ^7^. Subsequently, we demonstrated that the normal breasts of AA women contain elevated number of ZEB1+ cells that surround the ductal epithelial cells compared to EA women ^8^. Several different functions have been attributed to ZEB1 in breast cancer. For example, ZEB1-mediated epithelial to mesenchymal transition (EMT) confers a metastasis-initiating cell state to cancer cells ^9^. Stromal ZEB1 expression in breast cancer inversely correlates with abundance of multiple immune cell types ^10^. ZEB1 expression has been linked to claudin-low and metaplastic breast cancer subtypes, both of which are more common in AA compared to EA women ^11–14^. These observations prompted us to further characterize stromally located ZEB1^+^ cells for their potential role in breast cancer initiation and progression within the context of genetic ancestry.

In this study, we established several ZEB1^+^ cell lines, all derived from the breast tissues of AA women. These cells express PROCR and PDGFRα, similar to recently described stromal adipogenic progenitors cells of the mouse mammary gland that trans-differentiate into multiple cell types ^15^. Based on cell surface expression of PROCR and PDGFRα combined with ZEB1 expression, we have named these cells as ‘PZP cells’ for simplicity. Under appropriate growth conditions, these cells differentiated into adipogenic or osteogenic lineages and expressed the adipogenic marker Peroxisome Proliferator Activated Receptor gamma (PPARγ). Cell surface marker profiles revealed similarity of these cells to human prostate-derived mesenchymal stem cells (CD44^+^/CD90^+^/CD73^+^/CD105^+^ populations) ^16^. PZP:breast epithelial cell communication resulted in elevated expression of IL-6. Furthermore, epithelial cells acquired CD49f^+^/EpCAM^-^ basal cell characteristics, which has been shown to increase with aging and to increase susceptibility to breast cancer ^17, 18^. PZP cells, when transformed with oncogenes, generated metaplastic carcinomas in NSG mice suggesting that these cells are cell-of-origin of metaplastic carcinomas of the breast ^19^.

## MATERIALS AND METHODS

### Primary cell culture, Immortalization and Lentiviral transduction

Primary breast epithelial cell lines were created from fresh or cryopreserved, de-identified normal breast tissues donated to KTB by healthy AA women; all subjects provided written informed consent prior to donation. All experiments were carried out in accordance with the approved guidelines of the Indiana University Institutional Review Board. International Ethical Guidelines for Biomedical Research involving human subjects were followed. Generation of primary cells from cryopreserved tissues, culturing method and media composition have been described previously ^20^. While primary breast tissues cultured from EA donors generated predominantly tightly attached cuboidal epithelial cell colonies, tissues from AA donors yielded both loosely attached elongated cell clusters and cuboidal epithelial colonies. Differential trypsinization was used to separate two population of cells for immortalization. Cells were immortalized by human telomerase gene (hTERT) using the retrovirus vector pLXSN-hTERT or the lentivirus vector pCLX-PGK-hTERT vector (#114315, Addgene). Retrovirus and lentivirus preparations, infection of primary cells, and selection for immortalized cells by 100 µg/ml G418 (only with pLXSN-hTERT, 61-234, Corning) have been described previously ^21^. Cells were transformed with oncogenes *H-Ras^G12V^*, and SV40-T/t antigens expressing lentiviruses using vectors pLenti CMV-RasV12-Neo (w108-1) (HRAS ^G12V^, #22259, Addgene), and pLenti-CMV/TO-SV40 small + Large T (w612-1) (#22298, Addgene), respectively, as before ^22^. PZP KTB40 and KTB42 cells were modified to express Tomato Red or GFP markers using pCDH-EF1-Luc2-P2A-tdTomato (#72486, Addgene) or pCDH-EF1-Luc2-P2A-Cop GFP (#72485, Addgene) lentivirus. Cell lines in the laboratory are routinely tested for *Mycoplasma* ∼every 6 months (last testing was done on February 24, 2022).

### Genetic ancestry mapping and genotype analysis

Leukocyte DNA from the same donors was obtained from KTB. Blood was collected using the BD-Vacutainer spray-coated K3EDTA tubes (Becton Dickinson). Tubes were centrifuged for 15 min at 2000 RPM. Once the upper phase (plasma) was removed, the tubes were stored in -80°C. Peripheral blood leukocyte DNA was extracted using AutoGenprep 965 (Autogen, Inc.). Genotyping was performed using the KASP technology (LGC Genomics) and a 41-SNP panel (labeled 41-AIM panel) selected from Nievergelt *et al* ^23^. Genotype analysis using the 41 SNPs panel along with a Bayesian clustering method (Structure Software V2.3.4) was able to discern continental origins including European (Caucasian)/Middle East, East Asia, Central/South Asia, Africa, Americas, and Oceania. A reference set was obtained from the Human Genome Diversity Project (HGDP).

### Flow cytometry analysis and cell sorting

Adherent cells were collected by trypsinization, stained using antibodies PROCR (CD201)-PE (130-105-256), EpCAM-APC (130-091-254) EpCAM-PE (130-091-253), (Miltenyi Biotech Inc.), PDGFRα-PE (#323506, Biolegend), CD26-APC (#563670, BD Biosciences), CD105-PE (#12-1057-42, Invitrogen), CD73-PE (561014), CD90-APC (559869), CD44-APC (559942), CD24-PE (555428) (BD Pharmingen), CD10-APC (#340922, BD Biosciences), CD49f-APC (FAB13501A, R&D Systems), and were acquired using a BD LSR II flow cytometer. Data were analyzed using FlowJo software. Forward and side scatter were used to ensure that only live cells were considered in the analysis. Gating was done using appropriate FITC (555573), PE (555749) and APC (555576) (BD Pharmingen) isotype control antibodies and only a representative isotype control for two fluorescent markers are shown. Samples were analyzed and sorted using BD FACSAria and SORPAria.

### RNA extraction and quantitative real-time Reverse Transcription Polymerase Chain Reaction (qRT-PCR)

Total RNA was isolated using RNeasy Kit (74106, Qiagen) and 2 μg of RNA was used to synthesize cDNA with Bio-RAD iScript cDNA Synthesis Kit (170-8891, Bio-Rad). qRT-PCR was performed using Taqman universal PCR mix and predesigned gene expression assays with best coverage from Applied Biosystems. The following primer sets were used in our study:

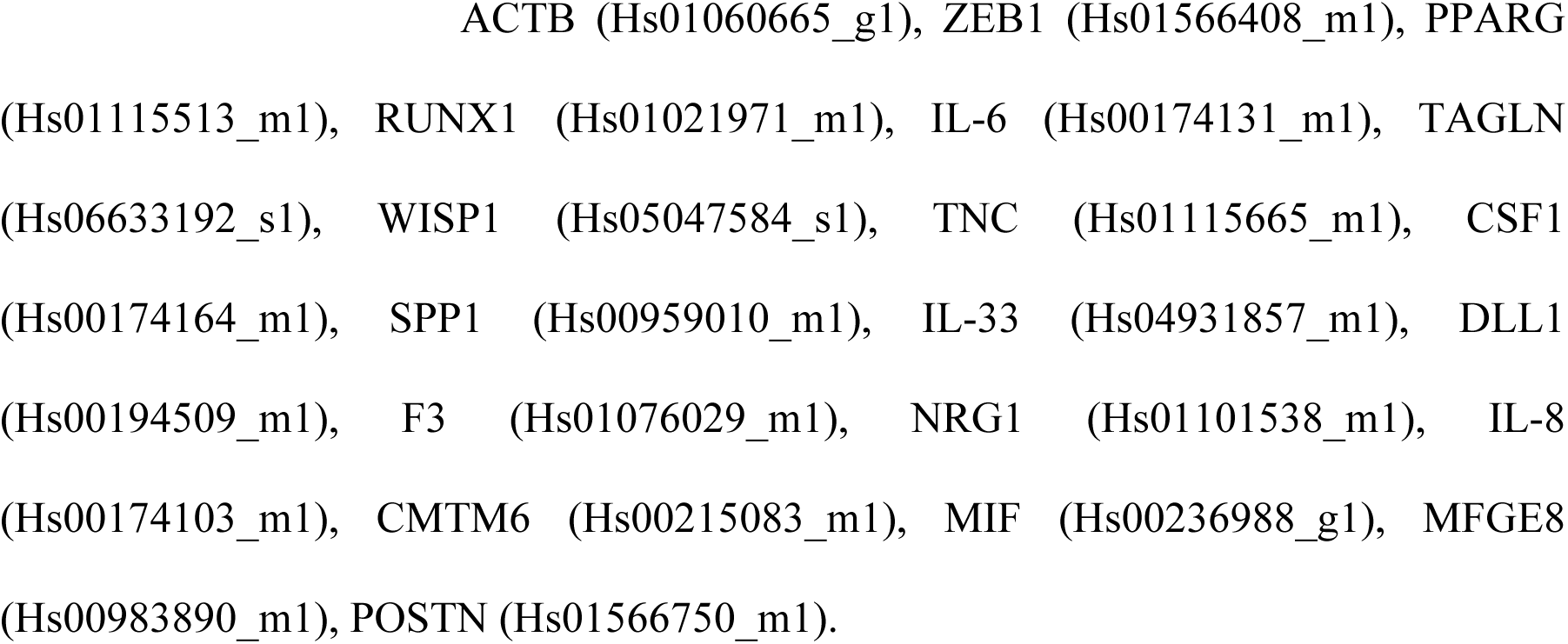

### Generation of TMA

We created a TMA comprising healthy breast tissue donated to the KTB (KTB-normal) from 49 women of African (AA), 154 of European (EA) and 46 of Hispanic/Latinas (LA) ancestry. The TMA also contained matched normal adjacent to tumor (NAT) and tumor tissue of approximately 50 donors. Part of the EA TMA has been described previously ^24^. Genetic ancestry information of KTB-normal donors and histopathology of tumors are shown in supplementary Figure 1 (Fig. S1) and Table S1, respectively. TMA was analyzed by immunohistochemistry (IHC) for PROCR (MAB22451, R&D Systems), ZEB1 (3G6, #14-9741-82, eBioscience), PDGFRα (#3174, Cell Signaling Technology), phospho-STAT3 (#ab30647, Abcam), Estrogen Receptor alpha (ERα, Clone EP1, Dako IR 084), GATA3 (sc-268, Santa Cruz Biotechnology), and FOXA1 (sc-6553, Santa Cruz Biotechnology) expression. IHC was done in a Clinical Laboratory Improvement Amendments (CLIA)-certified Indiana University Histopathology Laboratory and evaluated by three pathologists in a blinded manner. Quantitative measurements were done using the automated Aperio imaging system (ScanScope CS) and analysis was conducted using an FDA-approved algorithm. Positivity and H-scores were recorded and statistically analyzed as described previously ^8^. Data were analyzed in three different ways: (i) expression differences between AA, EA and LA KTB-normal; (ii) expression differences between KTB-normal and NATs; and (iii) expression differences between NATs and tumors.

**Table 1.** Compare H-score and Positivity between race in all patients within Normal tissue for PROCR, ZEB1 and PDGFRα.

| Marker | variable label | Race |  |  |  | P-value |
| --- | --- | --- | --- | --- | --- | --- |
|  |  | COLUMN_OVERALL | African American | European | Latino |  |
| PROCR | Positivity | 0.17 (0.04, 0.55) | 0.30 (0.06, 0.55) (N=31) | 0.15 (0.04, 0.41) (N=129) | 0.17 (0.04, 0.39) (N=33) | <.0001 |
|  | H-Score | 27.97 (5.45, 127.65) | 56.41 (10.46, 127.65) | 24.37 (6.68, 83.50) | 26.06 (5.45, 72.30) | <.0001 |
| ZEB1 | Positivity | 0.01 (0.00, 0.24) | 0.01 (0.00, 0.10) (N=33) | 0.01 (0.00, 0.12) (N=144) | 0.02 (0.00, 0.24) (N=28) | 0.0380 |
|  | H-Score | 1.62 (0.15, 30.72) | 2.21 (0.24, 16.47) | 1.46 (0.15, 19.76) | 2.75 (0.40, 30.72) | 0.0076 |
| PDGFRα | Positivity | 0.10 (0.01, 0.75) | 0.28 (0.03, 0.75) (N=35) | 0.09 (0.01, 0.73) (N=154) | 0.09 (0.02, 0.56) (N=18) | <.0001 |
|  | H-Score | 16.30 (1.43, 110.23) | 36.96 (6.25, 110.23) | 14.27 (1.43, 88.61) | 12.61 (2.69, 73.08) | <.0001 |

Note: Values expressed as median (min, max)
Note: P-value comparisons across race categories are based on Kruskal-Wallis

### Adipogenic and osteogenic differentiation assay

For adipogenic differentiation assay, PZP cells were plated in tissue culture dishes pre-coated with laminin-5-rich conditioned media from the 804G cell line in DMEM supplemented with 10% FBS (#26140-079) and 10 ng/ml bFGF (#3718-FB-010), and maintained at 37°C in a 5% CO2 incubator. Cells were allowed to grow to confluence and were then held at confluence for 2 days without changing of the media prior to exposure to the adipocyte differentiation cocktail (1 μg/ml insulin, 0.25 μg/ml dexamethasone, and 0.5 mM IBMX) in fresh media without addition of bFGF. The adipocytes differentiation media was changed every 3 days until day 12 for harvest. Cells were fixed and stained with oil red O (#ECM950, Millipore) according to the manufacturer’s instructions.

For osteogenic differentiation assay, PZP cells were plated onto tissue culture dishes in 50% DMEM/F-12 (#A4192001) supplemented with 10% FBS, 1% L-glutamine (#SCR028, Millipore Sigma). Cells were allowed to grow for 2-3 days to attain confluence, after which the normal media was removed and osteogenic differentiation medium was added. This media change corresponded to differentiation day 1. The osteogenic differentiation media contained 50% DMEM/F-12 supplemented with 10% FBS, 1% L-glutamine, 5 nM dexamethasone (#SCR028, Millipore Sigma), 250 μM L-ascorbic acid 2-phosphate (#SCR028, Millipore Sigma) and 10 mM β-glycerophosphate (#SCR028, Millipore Sigma). The media was changed twice a week. After 3 weeks, differentiated cells were washed in PBS and fixed in 95% methanol for 10 mins. Cells were stained with 2% alizarin red solution (Catalog no.# ECM815, Millipore) for 5 mins, then rinsed with water and imaged under fluorescence microscope to visualize the Ca^2+^ deposits.

### Co-culture and Cytokine/chemokine array

50% of each epithelial cell line (KTB34 and KTB39) and PZP cells (KTB40 and KTB42) were co-cultured in growth media for 2 days for RNA analysis. For cytokine/chemokine arrays, culturing of individual and co-culture was done as above and allowed to grow till the plates were confluent. Plates were washed with PBS and incubated with DMEM/F12 without other ingredients for 24 hours. Conditioned media (CM) was collected and subjected to cytokine/chemokine array. The cytokine/chemokine protein array was performed according to the manufacturer’s instructions (#ARY022B, R&D Systems).

### Antibodies and Western blotting

Primary antibodies used included rabbit anti-Ras (#3965, Cell Signaling Technologies), mouse anti-SV40-T antigens (MABF121, EMD Millipore), and mouse anti-β-actin (A5441, Sigma-Aldrich). Anti-mouse (#7076), and anti-rabbit (#7074) horseradish peroxidase (HRP) linked secondary antibodies were purchased from Cell Signaling Technologies. Cell lysates were prepared in radioimmunoassay buffer and analyzed by western blotting as described previously ^22^.

### Xenograft study and IHC of tumors and lungs

The Indiana University Animal Care and Use Committee approved the use of animals in this study and all procedures were performed as per NIH guidelines. Transformed cells with 50% basement membrane matrix (BME) type 3 (3632-005-02, Trevigen) (total 100 µL volume) were implanted into the mammary fat pad of 5 to 6 week old female NSG (NOD/SCID/IL2Rgnull) mice. A 60 day slow release 17β-estradiol (0.72 mg) pellet (SE-121, Innovative Research of America) was implanted at the time of mammary fat pad injection. Tumor growth was measured weekly and tumor volume was calculated as described previously ^22^. After 2-4 months, tumors and lungs were collected and processed for hematoxylin and eosin (H&E), ERα, GATA3, FOXA1, CK5/6 (IR 780, Dako), CK8 (35bH11, N1560, Dako), CK14 (LL002, ab7800, Abcam), and CK19 (IR 615, Dako) staining as described previously ^22^.

### Statistical analysis

Statistical analyses were conducted using Prism software program (version 6.0) and SAS software (version 9.4). Data were analyzed using one-way ANOVA. P values <0.05 were considered statistically significant. Nonparametric Wilcoxon rank-sum tests were used for unpaired analysis, as positivity and H-scores were not normally distributed, whereas nonparametric Wilcoxon signed-rank tests were used for paired analyses as described previously ^8^. For comparisons between three groups, the Kruskal-Wallis test was used.

## RESULTS

### Generation of <u>P</u>ROCR^+^/<u>Z</u>EB1^+^/<u>P</u>DGFRα^+^ (PZP) cell lines from breast tissues of healthy AA women

We created six immortalized PZP cell lines from AA (KTB40, KTB42, KTB32, KTB53, KTB57, and KTB59) by overexpressing human telomerase gene (hTERT) using primary cells isolated and propagated from core breast biopsies of self-reported healthy black women. Enrichment of African ancestry informative markers in these donors was confirmed through genotyping (**Fig. 1A**). To phenotypically characterize and document heterogeneity, these cells were subjected to flow cytometry using various epithelial, stem, mesenchymal, and fibroblast markers. These cell lines were predominantly PROCR^+^/EpCAM^-^ (**Fig. 1B, C, and** **Fig. S2A)**. To further characterize stem cell -related gene expression, we compared these immortalized variants with the immortalized PROCR^±^/EpCAM^+^ luminal/basal cell variants from EA and AA women we described previously ^21^. PROCR^+^/EpCAM^-^ cell lines expressed significantly higher levels of ZEB1 compared to PROCR^±^/EpCAM^+^ (KTB34 and KTB39) breast epithelial cell lines described previously ^21^ (**Fig. 1D**). Gene expression pattern in KTB34 cell line from EA overlaps Luminal-A intrinsic subtype of breast cancer, whereas gene expression in KTB39 from AA overlaps with Normal-like intrinsic subtype of breast cancer ^21^. Morphologically, PROCR^+^/EpCAM^-^ cell lines showed features of epithelial to mesenchymal transition (EMT), which is expected based on ZEB1 expression **(Fig. S2B)**.

**Figure 1:**
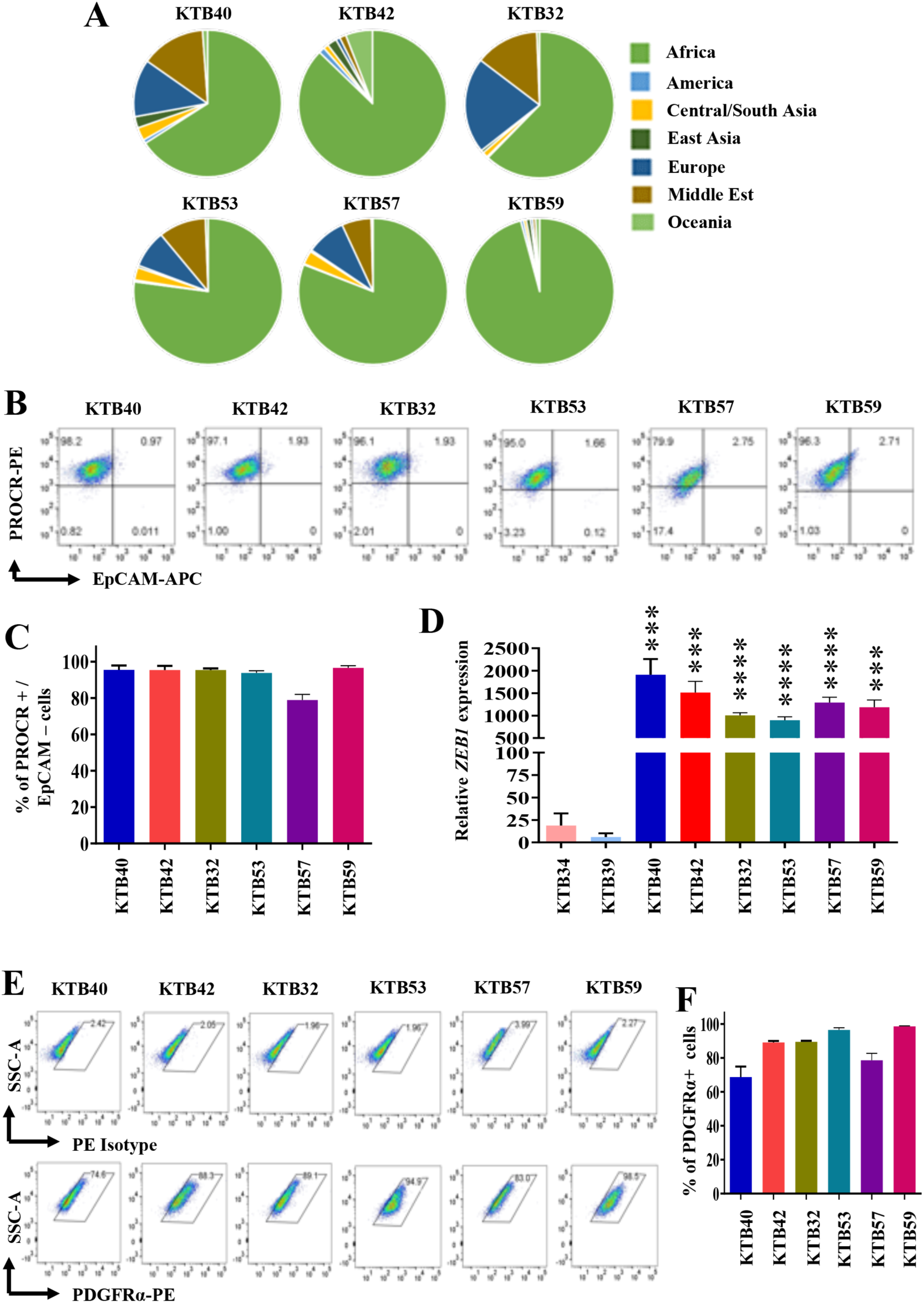
Establishment of PROCR^+^/ZEB1^+^/PDGFRα^+^ (PZP) cell lines from the healthy breast tissues of AA women. (A) Genetic ancestry mapping of breast tissue donors (KTB40, KTB42, KTB32, KTB53, KTB57, and KTB59) using a panel of 41-SNP. (B) PROCR^+^/EpCAM^‒^ cells are enriched in established cell lines from AA women. (C) Quantitation of PROCR^+^/EpCAM^‒^ cells. (D) ZEB1 expression levels in various PROCR^+^/EpCAM^‒^ cell lines compared to EpCAM^+^ (KTB34 and KTB39) luminal cell lines. (E) PROCR^+^/ZEB1^+^ cell lines express PDGFRα as determined by flow cytometry. (F) Quantitation of PDGFRα^+^ cells. ***p<0.001, ****p<0.0001 by ANOVA.

A recent study described PDGFRα^+^ stromal cells as adipogenic progenitors of the mammary gland that transdifferentiate into epithelial cells and migrate into the duct when stimulated by PDGF-C ^15^. Interestingly, these cells also expressed PROCR ^15^. We examined whether human breast PROCR^+^/ZEB1^+^ cells express PDGFRα. Indeed, >70% of cells were PDGFRα^+^ **(Fig. 1E and F; Fig. S2C)**, and these cells also expressed PDGFRβ **(Fig. S2D)**.

### Trans-differentiating properties of PZP cells

In order to determine to what extent PZP cells show similarity to adipogenic progenitors described in the mouse mammary gland and fibroadipogenic progenitors (FAPs) described in the skeletal muscle ^25^ with respect to trans-differentiation into adipogenic and osteogenic lineages, PZP cells were subjected to adipogenic and osteogenic differentiation growth conditions. Indeed, PZP cells differentiated into adipocytes (**Fig. 2A**) and the differentiated cells expressed adipocyte differentiation marker PPARγ (**Fig. 2B**). Adipogenic trans-differentiation capability of PZP cells was further confirmed through RUNX1 overexpression (**Fig. 2C**). PZP cells cultured in osteogenic media for 21 days showed osteogenic differentiation as evident from mineralization of matrix with Ca2^+^ and positive alizarin red staining (**Fig. 2D**). Thus, PZP cells enriched in the breasts of AA women could correspond to multipotent cells that can transdifferentiate into different cell types based on environmental cues.

**Figure 2:**
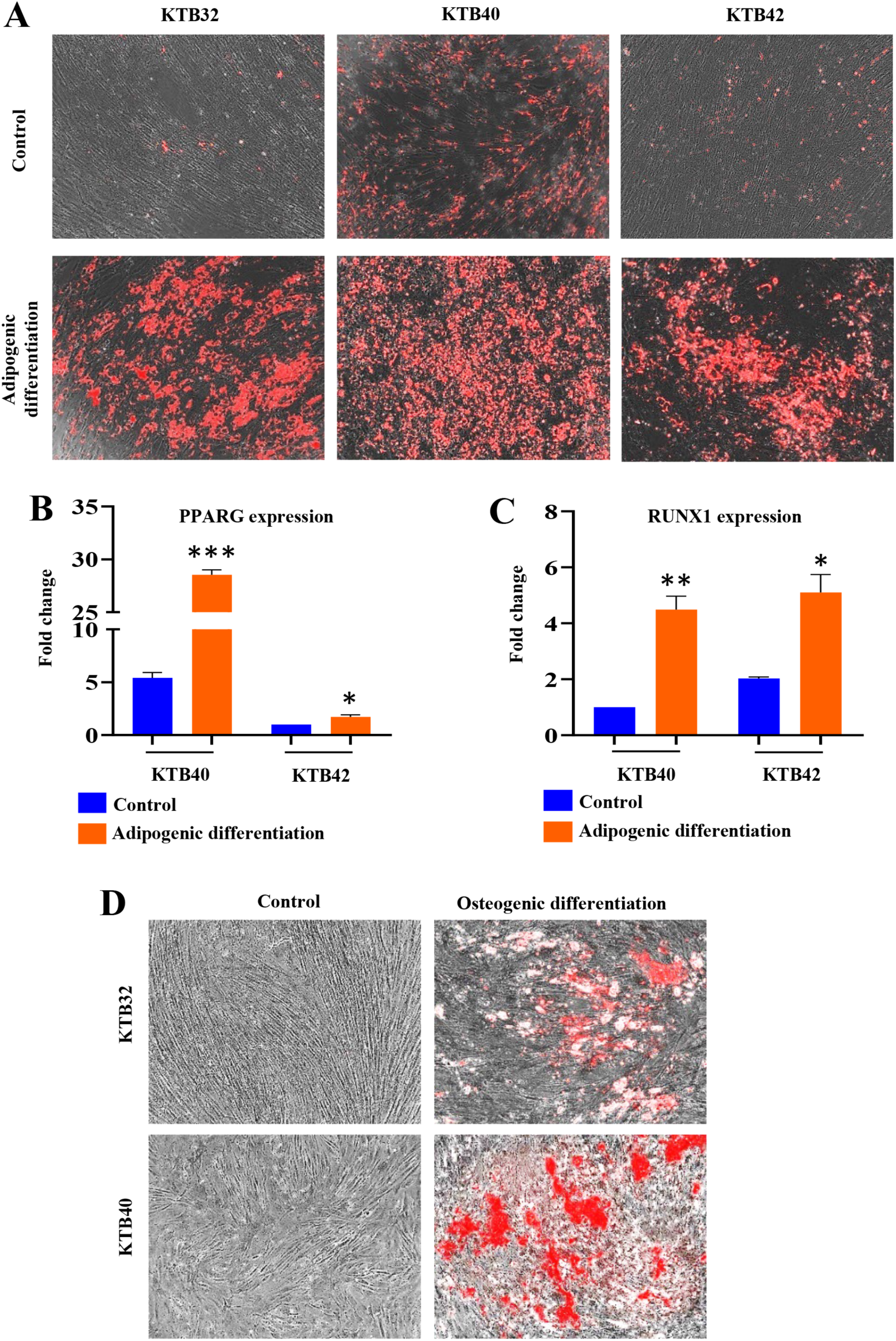
Trans-differentiating properties of PZP cells. (A) PZP cells undergo adipogenic differentiation under appropriate growth condition. Neural lipids stain red upon Oil Red-O staining. (B) Adipogenic differentiated PZP cells show elevated level of adipocytes differentiation marker PPARγ. (C) Adipogenic differentiation of PZP cells was confirmed through RUNX1 overexpression. (D) PZP cells undergo osteogenic differentiation under appropriate growth condition. Mineralization of matrix with Ca^2+^ was visualized by alizarin red staining. *p<0.05, **p<0.01, ***p<0.001 by ANOVA.

### PZP cells express lobular fibroblast markers

In breasts, loose connective tissue is unique to the terminal duct lobular units (TDLUs), which drain into the interlobular ducts, which in turn are embedded in a more dense connective tissue ^26^. CD105^high^ TDLU-resident lobular fibroblasts display properties different from interlobular fibroblasts ^27^. While the CD105^high^ lobular fibroblasts resemble mesenchymal stem cells (MSCs) both by phenotype and function, CD26^high^ interlobular cells remain fibroblast restricted ^27^. CD105^high^/CD26^low^ and CD105^low^/CD26^high^ lineages are considered to represent lobular and interlobular human breast fibroblastic cells (HBFCs), respectively ^28^. To further characterize the PZP cell lines, we examined the CD105 and CD26 staining pattern. PZP cells are enriched for CD105^high^/CD26^low^ population with inter-individual variability in the ratio between CD105^high^/CD26^‒^ and CD105^high^/CD26^low^ (**Fig. 3A - C**), which suggest that PZP cells reside predominantly in the lobules of breasts.

**Figure 3:**
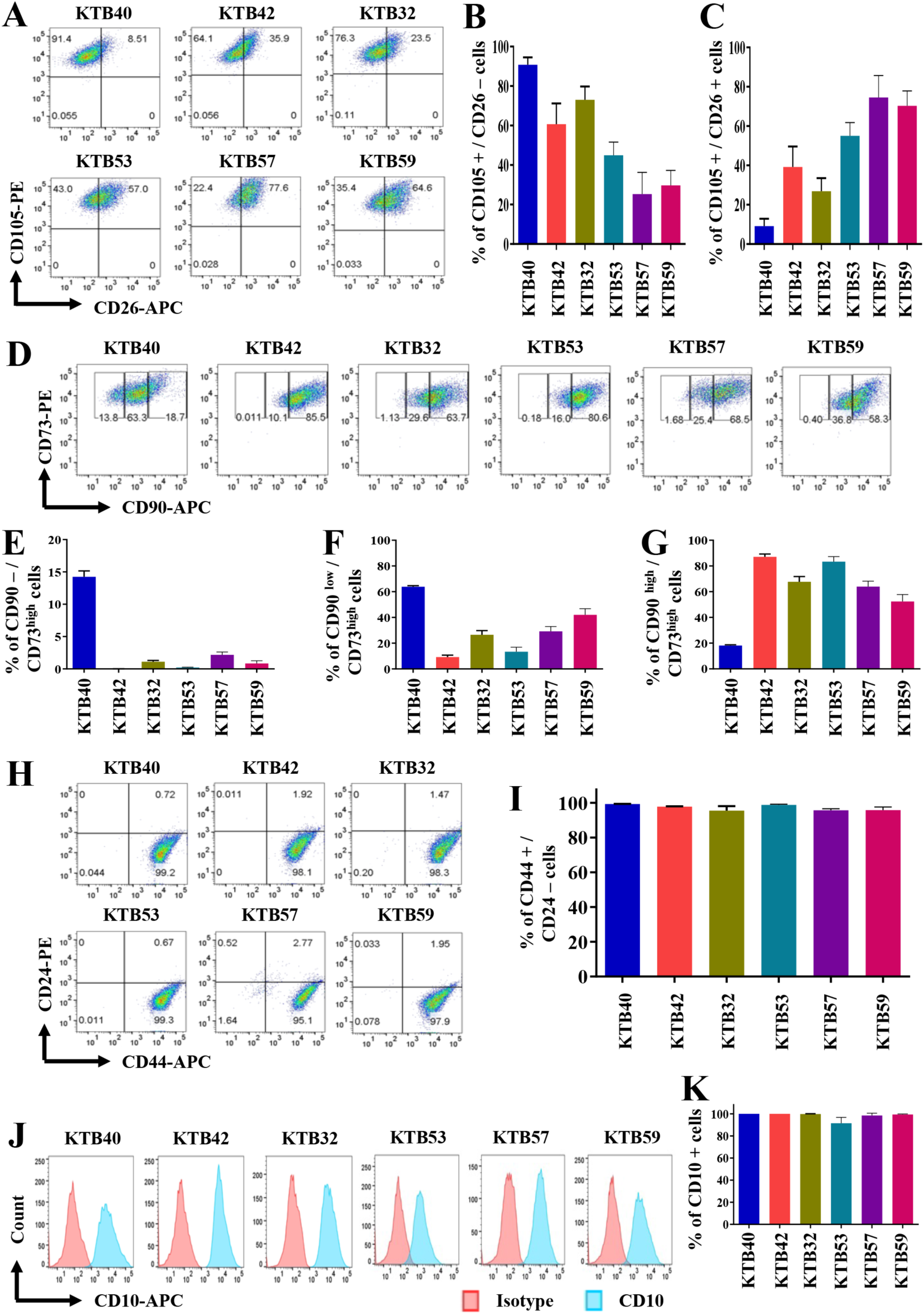
Phenotypic characterization of PZP cells. (A) PZP cell lines were stained with CD105 and CD26 antibodies to identify the lobular and interlobular origin of human breast fibroblastic cells. (B and C) Quantification of CD105^high^/CD26 ^̶^ and CD105^high^/CD26^low^ population of cells. (D) PZP cell lines were stained with CD90 and CD73 antibodies to identify rare endogenous pluripotent somatic stem cells and potential mesenchymal stem cells. (E, F and G) Quantification of CD90 ^̶^ /CD73^high^, CD90^low^/CD73^high^, and CD90^high^/CD73^high^ population of cells. (H) PZP cell lines were stained with CD44 and CD24 antibodies to determine whether their phenotype overlaps with cancer stem cells. (I) Quantification of CD44^+^/CD24^‒^ population. (J) PZP cell lines were stained with CD10 antibody to determine overlap with myoepithelial cell marker expression. (K) Quantification of CD10^+^ population.

CD90^‒^/CD73^+^ and CD73^+^/CD90^+^ are described as rare endogenous pluripotent somatic stem cells and potential mesenchymal stem cells, respectively ^29^. PZP cells contained CD90^‒^/CD73^+^ and CD73^+^/CD90^+^ subpopulations, with remarkable inter-individual variability in the ratio between CD90^‒^/CD73^+^, CD90^low^/CD73^+^ and CD90^high^/CD73^+^ (**Fig. 3D – G**). Isotype controls to generate gates are shown in **Fig. S2E.** CD44 and CD24 are the “original” markers used to characterize cancer stem cells (CSCs) in breast cancer ^30^. Interestingly, PZP cells displayed CD44^+^/CD24^‒^ phenotype (**Fig. 3H and I**). However, lack of EpCAM expression in PZP cells suggest that not all CD44^+^/CD24^-^ cells of the breast including breast tumors have stem cell properties. CD10 marker is used to isolate myoepithelial cells, although a recent study showed CD10 positivity of cancer associated fibroblasts ^31, 32^. PZP cells showed CD10^+^ phenotype (**Fig. 3J - K**). CD49f^high^/EpCAM^low^, CD49f^high^/EpCAM^medium^, and CD49f^low^/EpCAM^high^ cells are described as breast stem, luminal progenitor, and mature/differentiated cells, respectively ^33^. None of the PZP cell lines were positive for CD49f and EpCAM **(Fig. S2F)**.

### PZP cells are enriched in the normal breasts of women of AA compared to women of EA and Latina Ancestry (LA)

In our previous study, we had demonstrated that ZEB1+ cells in the normal breast are located in the stroma with close proximity to ductal epithelial cells ^8^. We generated a tissue microarray (TMA) comprising healthy breast tissues from 49 AA, 154 EA and 46 LA women. Ancestry marker distribution patten of donors is shown in **Fig. S1.** TMA was analyzed for protein levels of PROCR, ZEB1 and PDGFRα by IHC. Because of our previous observation of elevated ZEB1+ cells in the normal tissues adjacent to tumors (NATs) compared to breast tissues from healthy donors in case of EA ^8^, TMA also contained paired breast tumors and NATs of AA and EA women. To distinguish tissues of clinically healthy donors from NATs, clinically healthy tissues are labelled as KTB-normal. Pathologic features of tumors are described in **Table S1.** Representative IHC staining patterns of PROCR in KTB-normal, NATs, and tumors are shown in **Fig. 4A** and **B** and statistical analyses are presented in **Fig. 4C-E** and in **Table 1**. PROCR expression was observed in both ductal and stromal cells **(Fig. 4A** and **B).** PROCR-expressing cells were enriched in the KTB-normal breast tissue of AA women compared with EA and LA women (**Fig. 4B**, **Table 1)**. NATs of AA and EA women contained significantly higher levels of PROCR^+^ cells compared to KTB-normal suggesting the field effect of tumors on PROCR expression and NATs are not “normal” (**Fig. 4D**). Tumors of EA women displayed a modest increase in PROCR^+^ cells compared with those of NATs, while AA women did not show differences between NATs and tumors (**Fig. 4E**). Thus, PROCR^+^ cells are intrinsically higher in the normal breasts of AA women.

**Figure 4:**
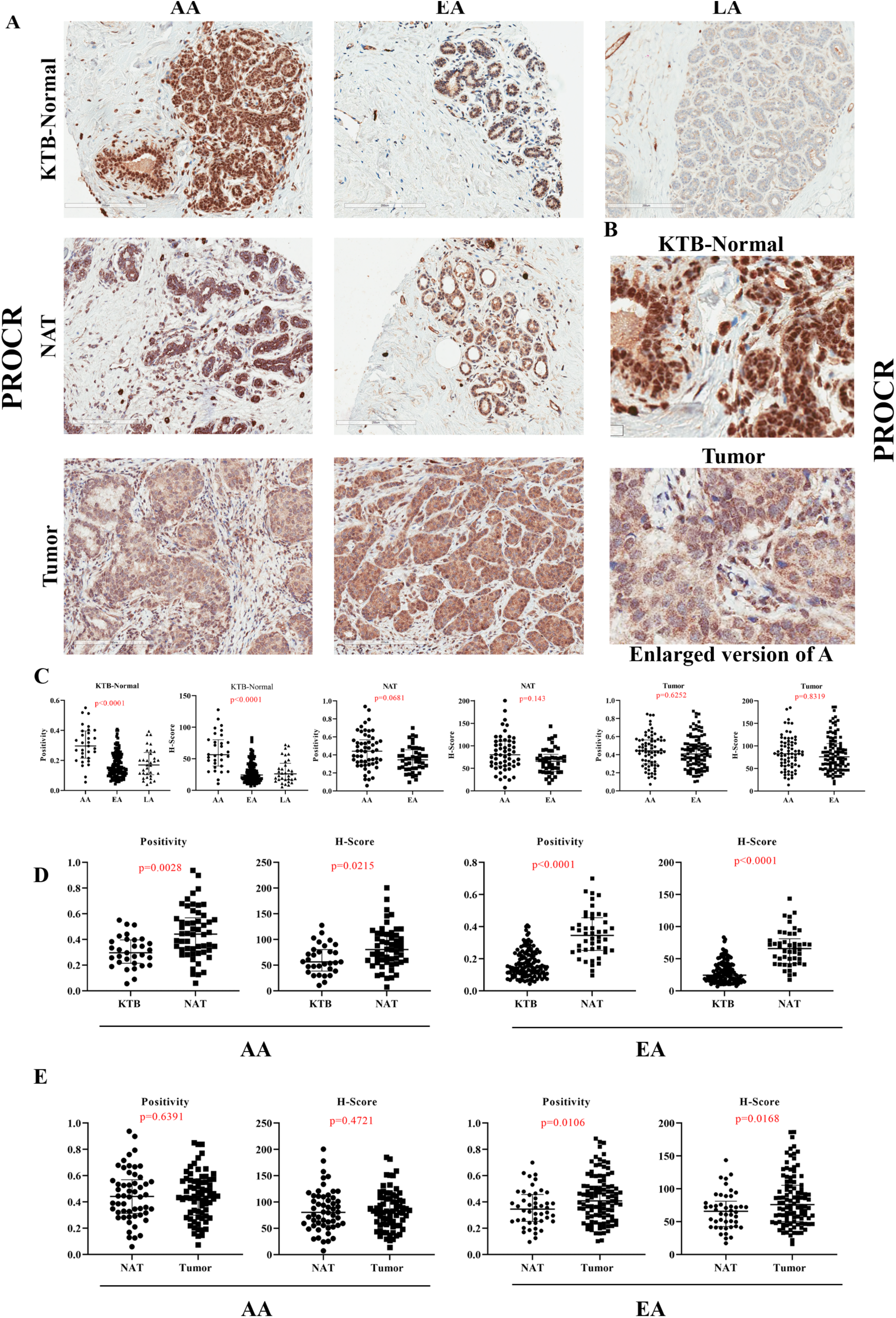
PROCR expression pattern in KTB-normal, NATs, and breast tumors. (A) Representative IHC of PROCR in KTB-normal, NATs, and/or tumors of AA, EA, and LA women. (B) Enlarged view of PROCR expression in KTB-normal and tumor. (C) Differences in PROCR expression (positivity and H-score) between KTB-normal of AA, EA and LA women. Differences in PROCR expression (positivity and H-score) between KTB-normal, NATs and/or tumors of AA, EA, and LA women. (D) Differences in PROCR expression (positivity and H-score) between KTB-normal and NATs in AA and EA women. (E) Differences in PROCR expression (positivity and H-score) between NATs and tumors in AA and EA women.

Consistent with our previous report regarding ZEB1 ^8^, KTB-normal breast tissues of AA women displayed higher levels of ZEB1^+^ cells compared to EA women. Interestingly, KTB-normal breast tissues of LA also demonstrated higher levels of ZEB1^+^ cells compared to EA women (**Fig. 5A-C**; **Table 1)**. Note remarkable inter-individual heterogeneity in ZEB1^+^ cells among LA women, possibly reflecting heterogeneity in genetic ancestry of LA. NATs of AA and EA women contained significantly higher levels of ZEB1^+^ cells compared to corresponding KTB-normal (**Fig. 5D**). Tumors of EA women displayed higher levels of ZEB1^+^ cells compared to NATs (**Fig. 5E**). Therefore, the normal breasts of AA women exhibit intrinsically higher levels of ZEB1^+^ cells but ZEB1+ cell numbers increase in the breast with cancer in EA women, consistent with our previous report ^8^.

**Figure 5:**
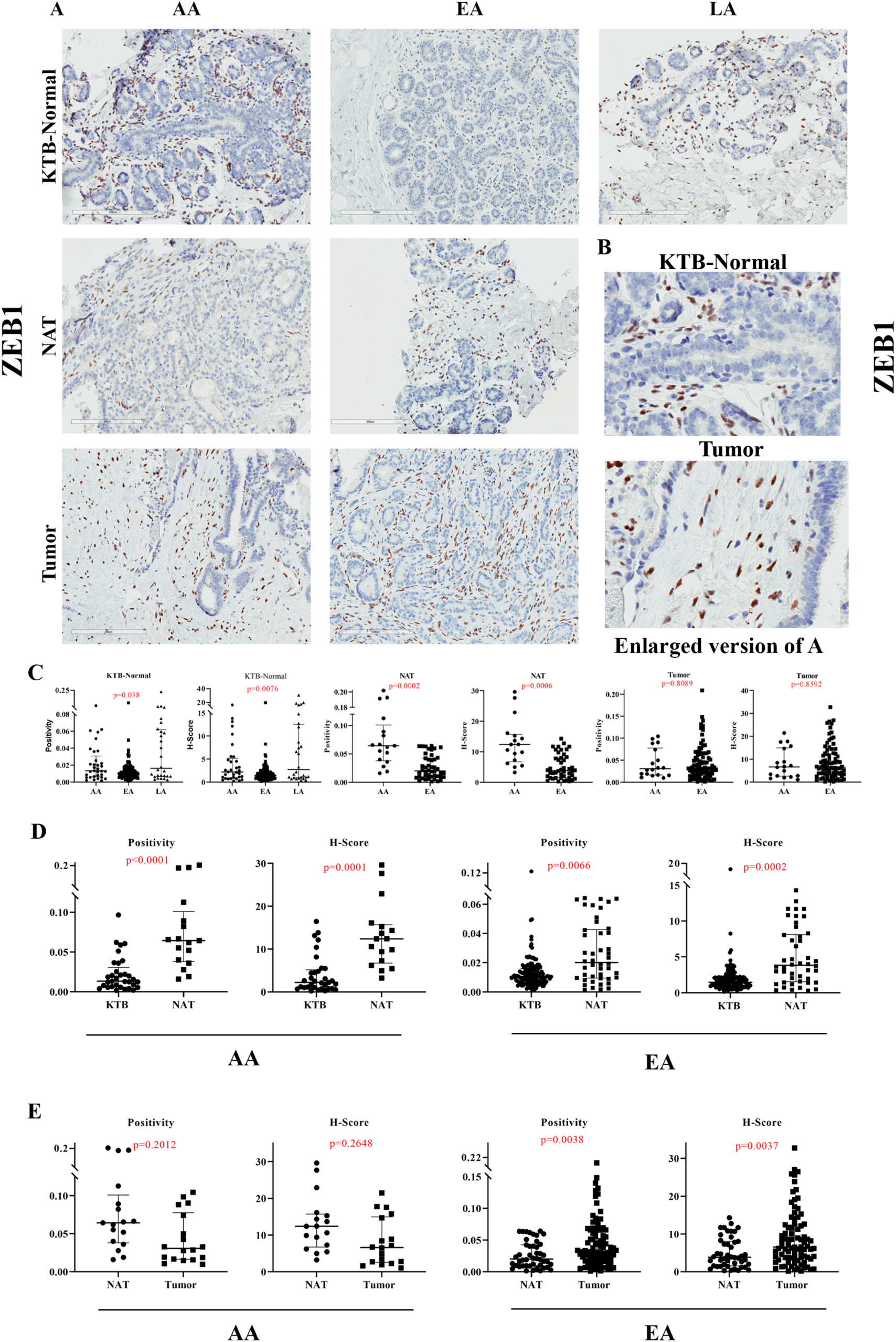
ZEB1 expression pattern in KTB-normal, NATs, and breast tumors. (A) Representative IHC of ZEB1 in KTB-normal, NATs, and/or tumors of AA, EA and LA women. (B) Enlarged view of ZEB1 expression in KTB-normal and tumor. Note ZEB1^+^ cells surround epithelial cell clusters. (C) Differences in ZEB1 expression (positivity and H-score) between KTB-normal of AA, EA and LA women. Differences in ZEB1 expression (positivity and H-score) between KTB-normal, NATs and/or tumors of AA, EA, and LA women. (D) Differences in ZEB1 expression (positivity and H-score) between KTB-normal and NATs in AA and EA women. (E) Differences in ZEB1 expression (positivity and H-score) between NATs and tumors in AA and EA women.

PDGFRα expression in the normal breast tissues was low and mostly in stromal cells (**Fig. 6A**). However, tumor epithelium showed some degree of positivity. PDGFRα-expressing cells were enriched specifically in the KTB-normal breast tissues of AA women compared with EA and LA women (**Fig. 6A-C**; **Table 1)**. Interestingly, KTB-normal breast tissues contained significantly higher levels of PDGFRα^+^ cells compared to NATs only in AA women, while no difference between KTB-normal breasts and NATs of EA women was observed (**Fig. 6D**). Furthermore, we did not observe a significant difference between NATs and tumors of AA women (**Fig. 6E**). Therefore, PDGFRα^+^ cells are intrinsically higher in the normal breasts of AA women. Taken together, our data suggest that PZP cells are elevated in the normal breasts of AA women compared to EA women, although expression of PROCR and PDGFRα in epithelial cells makes data interpretation bit difficult.

**Figure 6:**
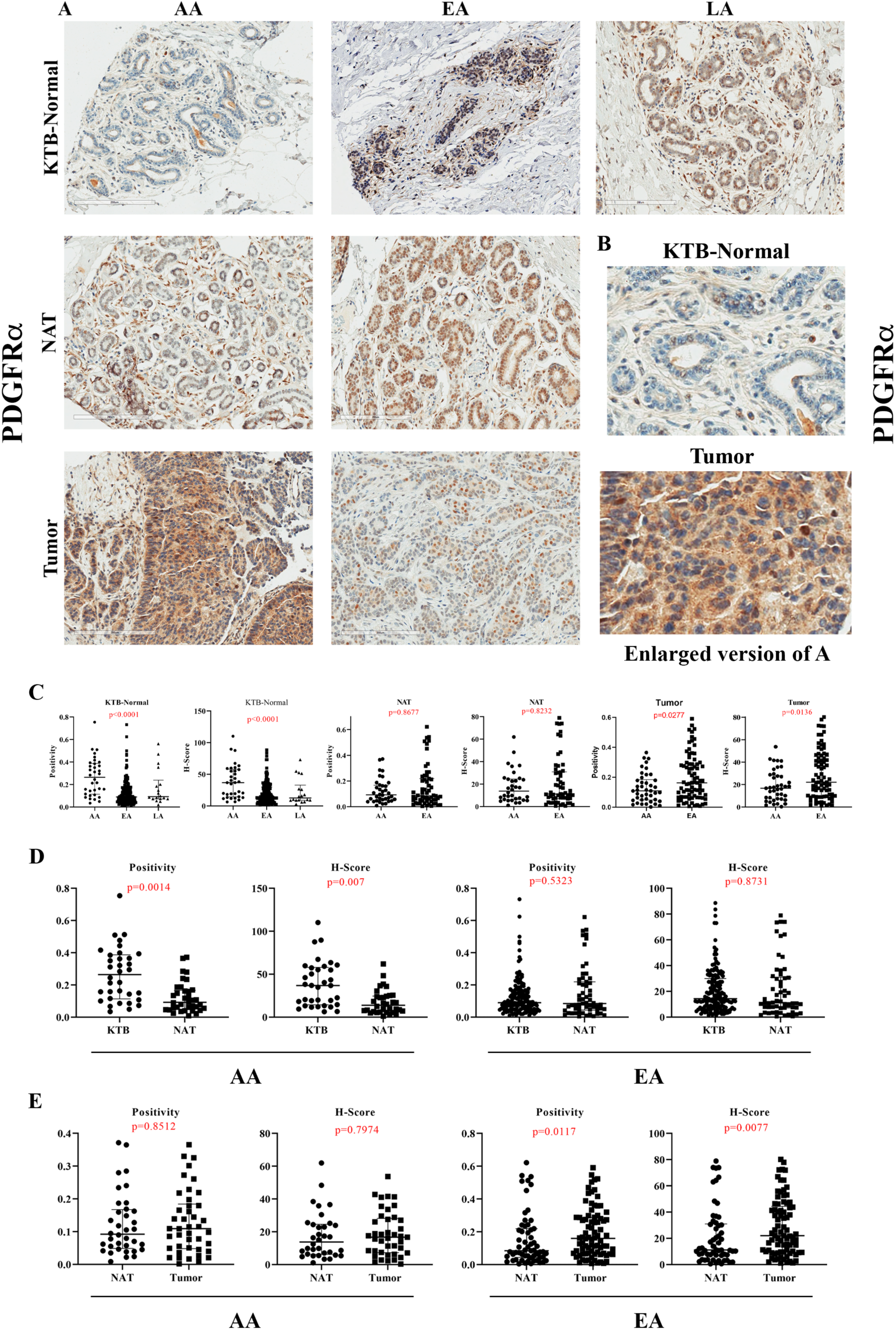
PDGFRα expression pattern in KTB-normal, NATs, and breast tumors. (A) Representative IHC of PDGFRα in KTB-normal, NATs, and/or tumors of AA, EA, and LA women. (B) Enlarged view of PDGFRα expression in KTB-normal and tumor. (C) Differences in PDGFRα expression (positivity and H-score) between KTB-normal of AA, EA, and LA women. Differences in PDGFRα expression (positivity and H-score) between KTB-normal, NATs and/or tumors of AA, EA, and LA women. (D) Differences in PDGFRα expression (positivity and H-score) between KTB-normal and NATs in AA and EA women. (E) Differences in PDGFRα expression (positivity and H-score) between NATs and tumors in AA and EA women.

We next examined the same TMA for luminal epithelial markers to rule out possible bias in our observations. Because FOXA1 along with another pioneer factor GATA3 and ERα form a lineage-restricted hormone-responsive signaling network in the normal breast^34^, we examined the expression levels of these markers. Representative staining patterns of ERα, GATA3 and FOXA1 are shown in **Fig. S3, Fig. S4** and **Fig. S5**, respectively. We did not observe genetic ancestry-dependent differences in ERα levels between the KTB-normal of AA, EA and LA women **(Fig. S3A-C)**. NATs of AA women contained modestly higher level of ERα^+^ cells compared with those of KTB-normal, while EA women did not show the difference **(Fig. S3D)**. Tumors of EA women contained significantly higher level of ERα^+^ cells compared with those of NATs, whereas AA women did not show the difference **(Fig. S3E),** which is consistent with pathologic features of tumors in EA as majority of them were ER^+^ tumors **(Table S1).** Similar to ERα, GATA3 expression in the normal breasts did not show genetic ancestry-dependent differences **(Fig. S4A-C)**. NATs of AA women contained significantly higher level of GATA3^+^ cells compared with those of KTB-normal, while EA women did not show the difference **(Fig. S4D)**. Tumors of EA women contained significantly higher level of GATA3^+^ cells compared with those of NATs, which is consistent with clinical features of tumors, whereas AA women did not show the difference **(Fig. S4E)**. FOXA1 also did not show any genetic ancestry-dependent differences between KTB-normal of AA, EA, and LA women **(Fig. S5A-C)**. NATs of AA and EA women contained significantly higher level of FOXA1^+^ cells compared with those of KTB-normal **(Fig. S5D)**. As expected, tumors of EA women contained significantly higher level of FOXA1^+^ cells compared with those of NATs, whereas AA women did not show the difference **(Fig. S5E)**. Importantly, these results suggest that the stromal but not the epithelial compartment of the normal breast shows genetic ancestry dependent variability in cell composition, at least with markers examined.

### Modeling the effects of PZP cells on breast tumorigenesis

To obtain potential insight into signaling pathway alterations in epithelial and PZP cells as a consequence of their cross-talk, we performed cytokine/chemokine profiling of factors secreted by an immortalized luminal epithelial cell line, a PZP cell line and both co-cultured together (50% of each cell line) for ∼24 hours. If the expression under the co-culture condition was much higher than expression in either cell types alone, we interpreted those results as cooperative effect on gene expression. If the expression under co-culture condition was the same or lower than expression in either cell type alone, we interpreted those results as no effect of cell-cell interaction. We selected the Luminal-A epithelial cell line KTB34 for this experiment because this cell line upon transformation generates adenocarcinoma, similar to human breast cancer ^21, 22^. While luminal epithelial cell line expressed several ligands such as PDGF-AA and osteopontin, which can affect trans-differentiation of PZP cells, PZP cells expressed factors such as EGF, HGF and SDF-1α, which can signal in luminal cells **(Fig. 7A; Table 2)**. Interestingly, IL-6 is produced only under coculture condition **(Fig. 7A; Table 2)**. We further confirmed the IL-6 production under co-culture condition at mRNA level by qRT-PCR (**Fig. 7B**). We suspect PZP cells produce IL-6 in response to interaction with luminal cells as basal expression of IL-6 was much higher in PZP cells compared to luminal cell line **(Fig. S6A)** and luminal cells secreted IL-1α, which we have previously shown to induce IL-6 in stromal cells ^35^. There appears to be specificity in cytokine production under co-culture conditions as we did not observe elevated production of IL-8 under co-culture condition of PZP and epithelial cells (**Fig. S6B**).

**Figure 7:**
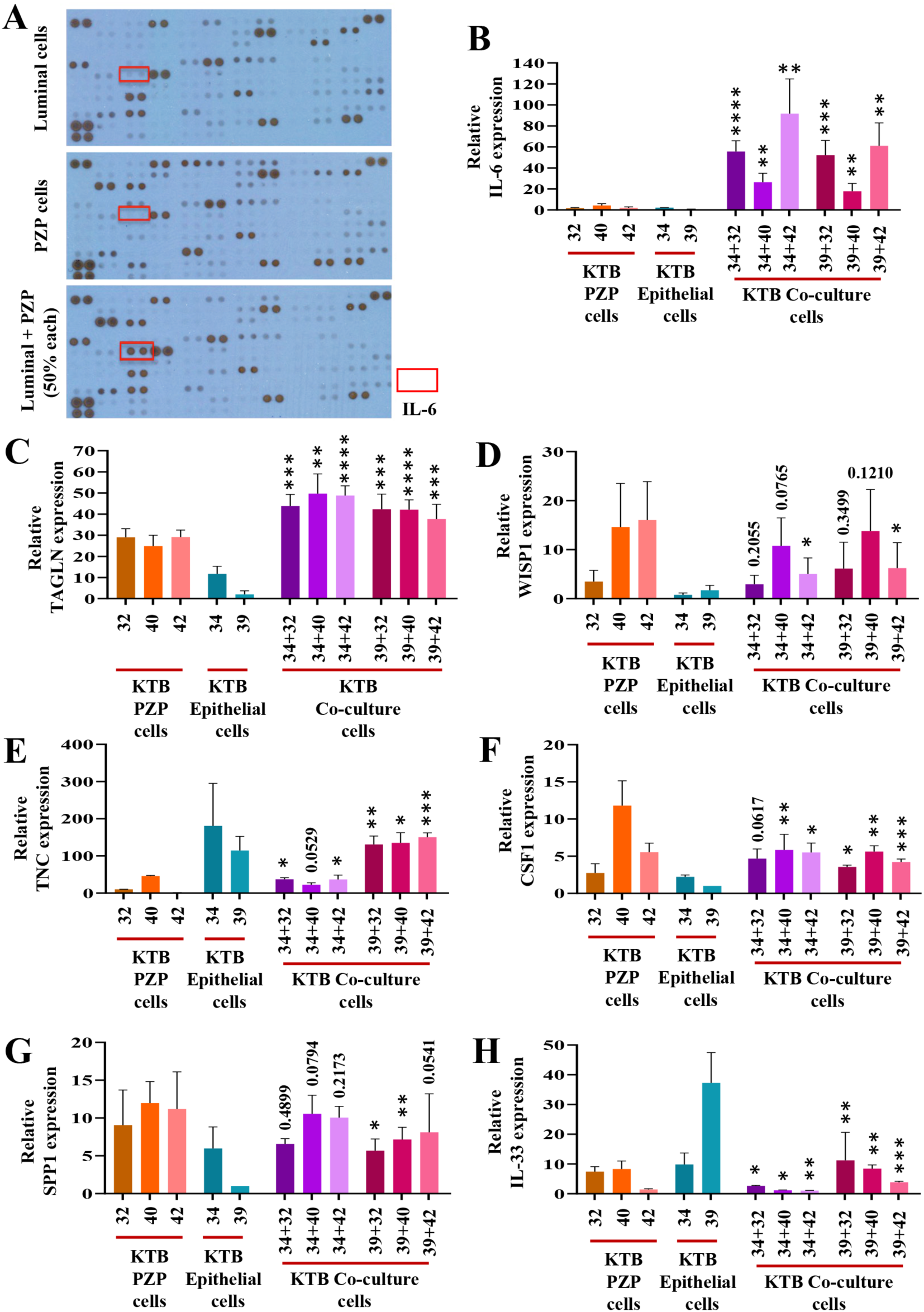
PZP-Epithelial cells interaction alters the gene expression. (A) IL-6 expression was detected only when luminal and PZP cells were co-cultured. R&D systems cytokine/chemokine array was used to identify secreted factors by luminal and PZP cells either alone or together (50% each). Expression of IL-6 (B), TAGLN (C), WISP1 (D), TNC (E), CSF1 (F), SPP1 (G), and IL-33 (H) in PZP cell lines (KTB32, KTB40, KTB42), luminal progenitor (epithelial cells; KTB34, KTB39), and co-culture of PZP and luminal progenitor cell lines. Statistical significance (*p* values) was determined by comparing KTB32/40/42 (PZP cells), KTB34/KTB39 (epithelial cells) and respective co-cultured PZP + epithelial cells as indicated in the figure. *p<0.05, **p<0.01, ***p<0.001, ****p<0.0001 by ANOVA.

**Table 2.**
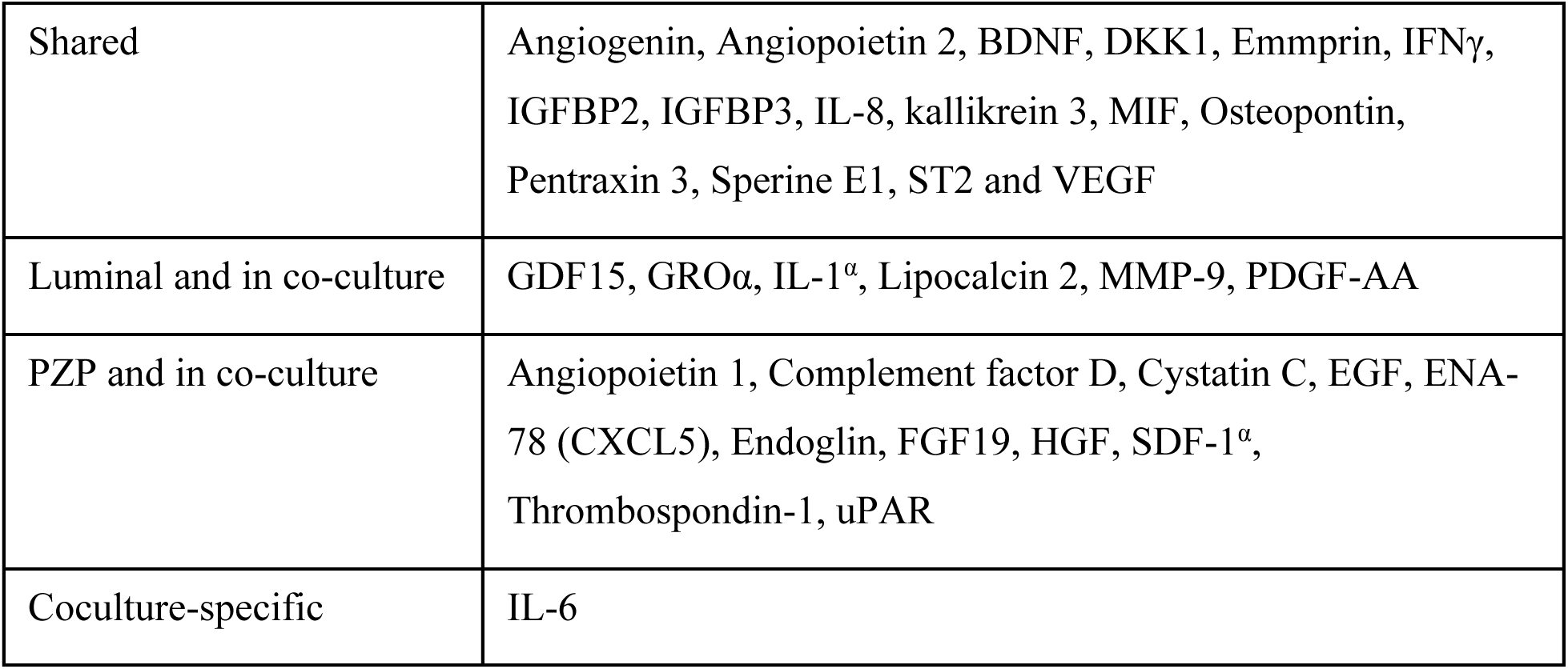
Cytokine/chemokine array identified factors secreted by luminal and PZP cells either alone or together.

|  |  |
| --- | --- |
| Shared | Angiogenin, Angiopoietin 2, BDNF, DKK1, Emmprin, IFNγ, IGFBP2, IGFBP3, IL-8, kallikrein 3, MIF, Osteopontin, Pentraxin 3, Sperine E1, ST2 and VEGF |
| Luminal and in co-culture | GDF15, GROα, IL-1α, Lipocalcin 2, MMP-9, PDGF-AA |
| PZP and in co-culture | Angiopoietin 1, Complement factor D, Cystatin C, EGF, ENA-78 (CXCL5), Endoglin, FGF19, HGF, SDF-1α, Thrombospondin-1, uPAR |
| Coculture-specific | IL-6 |

### PZP-epithelial cells interaction alters the expression of specific genes

We performed an extensive literature search to identify potential genes whose expression could be altered due to stromal-epithelial cell interaction. For example, transgelin (TAGLN) is known to be a specific marker of smooth muscle differentiation and highly expressed in the myoepithelial cells and fibroblastic cells of benign breast tissue with limited expression in luminal cells ^36, 37^. A population of subepithelial cells that lines the entire villus-crypt axis of intestine express high levels of PDGFRα, DLL1, F3, and EGF-family ligand Neuregulin 1 (NRG1) ^38^. In addition, NRG1 is also expressed in mesenchymal cells adjacent to the proliferative crypts ^38^. Expression of these genes were examined in individual cell types and under co-culture. Three PZP cell lines (KTB32, KTB40, KTB42) and two epithelial cell lines (KTB34, KTB39), and co-culture of PZP and epithelial cell lines (50% of each cell line) were used. We observed abundant TAGLN expression in PZP cells, while epithelial cells expressed at low level. Interestingly, the expression of TAGLN was further increased under co-culture condition (**Fig. 7C**). We found a low level of DLL1, F3, and NRG1 expression in PZP cell lines except high level of NRG1 in KTB42. DLL1, F3 and NRG1 are expressed predominantly in epithelial cell lines. In co-cultured cells, expression of DLL1 and F3 was additive depending on the cell type **(Fig. S6C - E)**. Taken together, these results indicate that PZP cells correspond to stromal cells that interact with epithelial cells to alter gene expression in a reciprocal manner. However, these cells are unlikely to function similar to subepithelial mesenchymal cells described in the intestine ^38^.

### Potential role of PZP cells in immune cell modulation in the microenvironment

Apart from IL-6, several other factors have been shown to alter the immune microenvironment. For example, WNT1-inducible-signaling pathway protein 1 (WISP1) expression affects the clinical prognosis through associations with macrophage M2 polarization, and immune cell infiltration in pan-cancer and helps to maintain CSC properties in glioblastoma ^39^. PZP cell lines displayed higher expression of WISP1 compared to luminal cell lines, while additive expression was observed under co-culture (**Fig. 7D**). Tenascin-C (TNC) promotes inflammatory response by inducing the expression of multiple proinflammatory factors in innate immune cells such as microglia and macrophages. TNC drives macrophage differentiation and polarization predominantly towards an M1-like phenotype ^40^. TNC is expressed mostly by epithelial cells and this expression was unaffected under co-culture (**Fig. 7E**). CSF1 controls both the differentiation and immune regulatory function macrophages ^41^. PZP cell lines displayed higher expression of CSF1 compared to epithelial cell lines and the expression unaffected under co-culture (**Fig. 7F**). Osteopontin (SPP1 or OPN), secreted by myofibroblasts, promotes M2 macrophage polarization through STAT3/PPARγ pathway ^42^. PZP cell lines displayed high expression of SPP1 compared to epithelial cell lines and the expression remained additive under co-culture condition (**Fig. 7G**). IL-33 is known to be upregulated in metastases-associated fibroblasts, and the upregulation of IL-33 activates type 2 inflammation in the metastatic microenvironment and facilitates eosinophils, neutrophils, and inflammatory monocytes recruitment to lung metastases ^43^. Co-culturing of PZP and epithelial cells did not affect IL-33 expression (**Fig. 7H**). CMTM6 maintains the expression of PD-L1 in tumor cells to regulate anti-tumor immunity ^44^. Epithelial cells but not PZP cells expressed higher levels of CMTM6, which was unaffected by co-culture condition **(Fig. S6F)**. Macrophage migration inhibitory factor (MIF) is an essential cytokine that is involved in the regulation of macrophage function in host defense through the suppression of anti-inflammatory effects of glucocorticoids ^45^. Both PZP and epithelial cell lines expressed MIF, but co-culture condition did not further effect expression **(Fig. S6G)**. Secretion of MFGE8 can reprogram macrophages from an M1 (proinflammatory) to an M2 (anti-inflammatory but pro-tumorigenic) phenotype ^46^. MFGE8 also induces the production of basic fibroblast growth factor that is responsible for fibroblast migration and proliferation ^47^. PZP cell lines displayed high expression of MFGE8 which was unaffected by co-culture **(Fig. S6H)**. Periostin (POSTN) is predominantly secreted by stromal fibroblasts to promote the proliferation of tumor cells. POSTN is also an essential factor for macrophage recruitment in the tumor microenvironment and involved in the interactions between macrophages and cancer cells ^48^. PZP cell lines displayed high expression of POSTN and the expression was unaffected by co-culture condition **(Fig. S6I)**. We noted variation in the expression of select genes amongst PZP cell lines (WISP1, CSF1, NRG1, and POSTN, for example), suggesting inter-individual variability in gene expression in stromal cells, similar to epithelial cell types we described previously ^21^. Collectively, these results suggest that PZP-epithelial interaction in the breast could impact the levels of select chemokines/cytokines in the breast environment (IL-6 and TAGLN for example) with consequential effects on the normal and/or tumor immune environment.

### The effects of breast cancer: PZP cell interaction on trans-differentiation of epithelial luminal progenitor cells

In order to further investigate the intercellular communication between PZP and epithelial cells, we generated stable tdTomato-labeled KTB40 and KTB42 cell lines using pCDH-EF1-Luc2-P2A-tdTomato lentivirus **(Fig. S7A)**. To determine whether co-culturing alter the phenotype of epithelial cells, tdTomato-labeled PZP (KTB40, KTB42), epithelial (KTB34, KTB39), and co-cultured PZP and epithelial cells were analyzed by flow cytometry using CD49f and EpCAM, which can differentiate luminal mature (CD49f^-^/EpCAM^+^), luminal progenitor (CD49F^+^/EpCAM^+^) and basal/stem (CD49f^+^/EpCAM^-^) cells. Co-cultured epithelial cells displayed an increase in CD49f^+^/EpCAM^-/low^ and reduced CD49f^-/low^/EpCAM^+/high^ cell populations in co-culture condition **(Fig. 8A-C; Fig. S7B)**. tdTomato-Red+ PZP cells on their own were CD49f^-^/EpCAM^-^), a small fraction of these cells displayed a unique CD49f^+^/EpCAM^+^ phenotype under co-culture condition **(Fig. 8D and E; Fig. S7C)**. These results suggest that similar to mouse mammary fibroadipogenic cells ^15^, PZP cells can potentially acquire epithelial characteristics under specific conditions. Our attempts of characterize flow-sorted CD49f^+^/EpCAM^+^ trans-differentiated PZP cells were not successful because of dominant growth of few contaminating PZP cells in the sorted population of cells.

**Figure 8:**
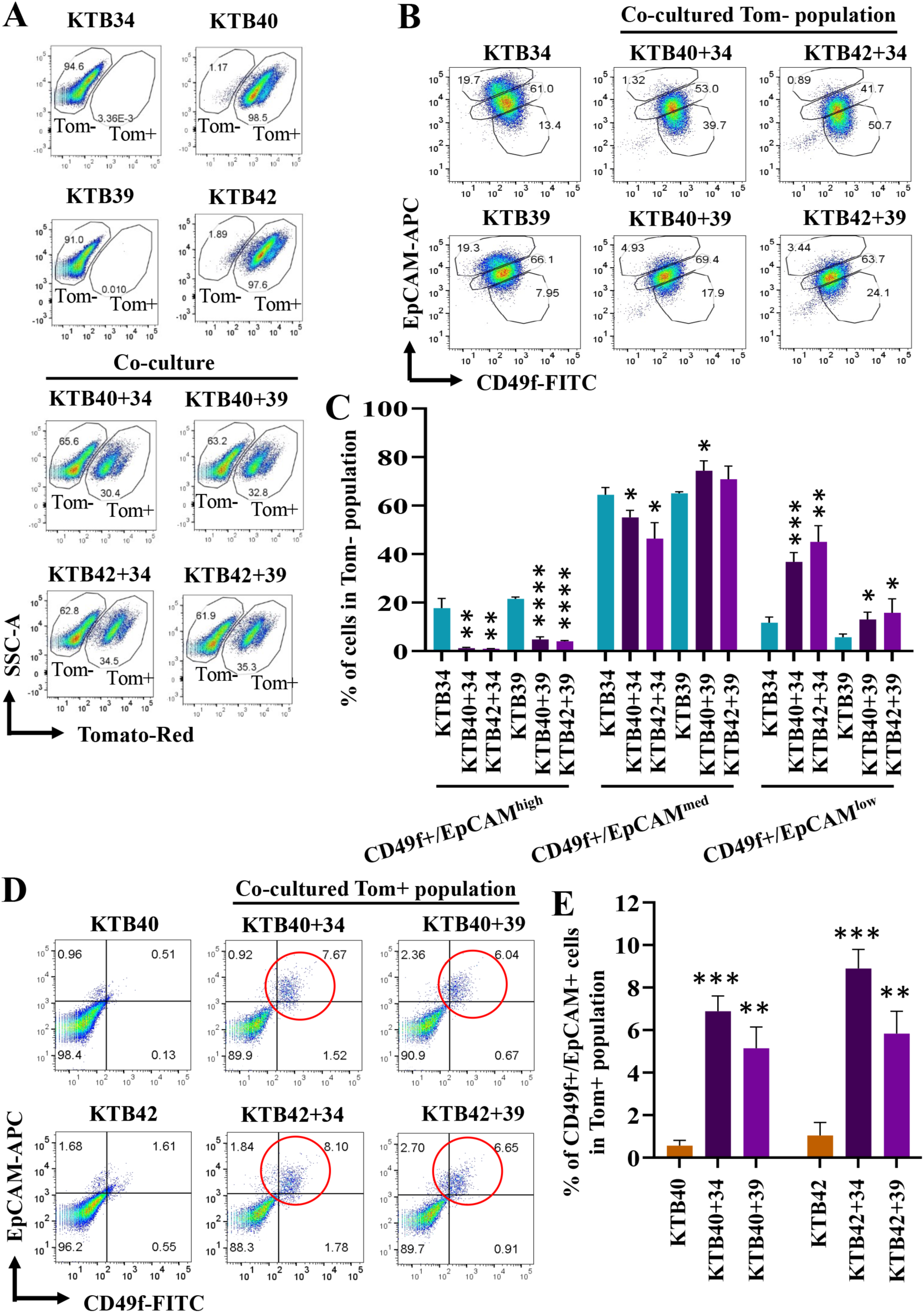
The effects of PZP cells on trans-differentiation of epithelial luminal progenitor cells. (A) Gating of tomato- (Tom-) and tomato+ (Tom+) populations of KTB34, KTB39, and co-cultured KTB40/KTB42 and KTB34/KTB39 cell lines. (B) CD49f and EpCAM staining patterns of KTB34, KTB39, and co-cultured KTB40/KTB42 and KTB34/KTB39 cell lines. (C) Quantification of CD49f^+^/EpCAM^high^, CD49f^+^/EpCAM^med^, and CD49f^+^/EpCAM^low^ populations in Tom-cell population. (D) CD49f and EpCAM staining patterns of KTB40, KTB42, and co-cultured KTB40/KTB42 and KTB34/KTB39 cell lines. Only Tom+ population was analyzed. (E) Quantification of CD49f^+^/EpCAM^+^ population among Tom+ population. *p<0.05, **p<0.01, ***p<0.001, ****p<0.0001 by ANOVA.

### Transformed PZP cells generate metaplastic carcinoma

We recently reported that cell-of-origin but not the oncogenic mutations determine the histotypes of breast tumors using breast epithelial cell lines derived from multiple donors and a defined set of oncogenes ^22^. We used the same strategy to determine whether PZP cells are cell-of-origin of specific malignancy of the breast. Mutant Ras is one of the potent oncogenes used to transform breast epithelial cells *in vitro* ^49^. Although initially considered not a relevant oncogene in breast cancer and breast cancer-related studies that utilized Ras oncogenes were often dismissed upon by reviewers, recent studies have clearly shown the role of Ras oncogene in endocrine resistance and metastasis of luminal breast cancer ^50^. Furthermore, the majority of breast cancer cell lines used as model systems have either mutation in Ras genes themselves or in downstream effectors or negative regulators of Ras pathway and Ras pathway is the major signaling pathway activated in TNBCs and Claudin-low breast cancer subtype ^51^. We transformed PZP cell lines with HRas^G12V^, SV40-T/t antigen and combination of both HRas^G12V^ and SV40-T/t antigen using lentivirus, since this combination was the most effective in breast epithelial cell transformation ^22^. Western blotting was used to detect the overexpression of mutant HRas^G12V^ **(Fig. S8A)**, SV40-T/t antigen **(Fig. S8B)**, and combination of both HRas^G12V^ and SV40-T/t **(Fig. S8C)**. Phase contrast images of PZP transformed cell lines are shown in **Fig. S8D**. Transformation of PZP cells with activated HRas^G12V^ increased the fraction of cells that have acquired epithelial phenotype and express EpCAM, particularly in KTB42 **(Fig. 9A and B; Fig. S8D)**. Transformed PZP cells expressed the stem/basal cell marker CD49f with inter-individual variability **(Fig. 9A; Fig. S9A)**. Transformation also altered the cell surface profiles of mesenchymal stem cell marker CD90 **(Fig. 9C; Fig. S9A)**. Transformed PZP cells were CD201^+^ and CD44^+^ **(Fig. 9B; Fig. S9A and S9B)**. Thus, PZP cells transdifferentiate upon transformation by acquiring CD49f expression.

**Figure 9:**
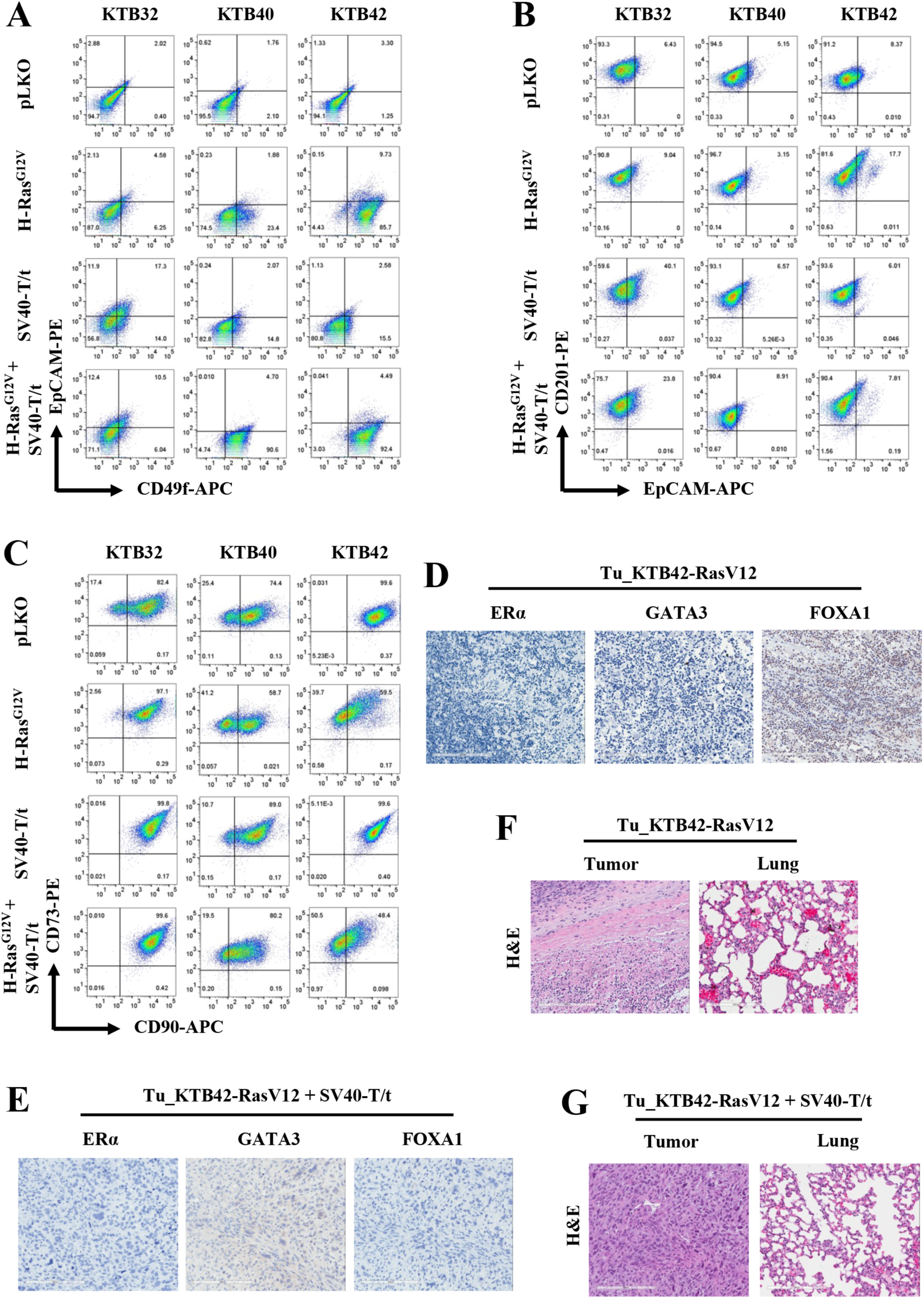
PZP cells transformed with Ras and SV40-T/t antigen are tumorigenic. (A) CD49f and EpCAM staining patterns of immortalized and transformed PZP (KTB32, KTB40 and KTB42) cell lines. CD49f^+^/EpCAM ^̶^, CD49f^+^/EpCAM^+^ and CD49f ^̶^ /EpCAM^+^cells correspond to stem/basal, progenitor and differentiated cells, respectively. (B) CD201 and EpCAM staining patterns of immortalized and transformed PZP (KTB32, KTB40, and KTB42) cell lines. (C) CD90 and CD73 staining patterns of immortalized and transformed PZP (KTB32, KTB40, and KTB42) cell lines. (D) IHC analyses of luminal markers ERα, GATA3 and FOXA1 in tumors developed from KTB42-HRas^G12V^ transformed cells. (E) IHC analyses of luminal markers ERα, GATA3 and FOXA1 in tumors developed from the KTB42-HRas^G12V^+SV40-T/t antigen transformed cells. (F) Tumor developed from KTB42-HRas^G12V^ transformed cells did not show extensive lung metastasis. (G) Tumor developed from KTB42-HRas^G12V^+SV40-T/t antigen transformed cells did not show lung metastasis. H&E staining shows the type of tumors.

We next determined whether cells expressing oncogenes are tumorigenic in NSG mice. Indeed, five million transformed cells in 50% matrigel implanted into the mammary gland of 6-7 week old female NSG (NOD/SCID/IL2Rgnull) mice (n=5) progressed into tumors. All animals injected with HRas^G12V^ + SV40-T/t and four out of five animals injected with HRas^G12V^ transformed cells but not parental immortalized cell lines (n=5) generated tumors. Tumor was resected and analyzed by H&E staining and expression of luminal markers estrogen receptor alpha (ERα), GATA3, and FOXA1 and cytokeratins CK5/6, CK8, CK14 and CK19. The luminal cells express cytokeratin 19 (CK19), while basal cell types express cytokeratin 5/6 (CK5/6) and cytokeratin 14 (CK14), and cells expressing both CK14 and CK19 show luminal progenitor phenotype or luminal/basal hybrid cells of the breast ^52^. KTB42-HRas^G12V^ cell-derived tumor was ERα^-^/GATA3^-^ /FOXA1^-^ (**Fig. 9D**). Surprisingly, KTB42-HRas^G12V^-derived tumor was also CK5/6^-^/CK8^-^/CK14^-^/CK19^-^ **(Fig. S9C)**. KTB42 cell line transformed with both mutant HRas^G12V^ and SV40-T/t antigen also developed tumors in NSG mice. KTB42-HRas^G12V^+SV40-T/t cell-derived tumor was ERα^-^/GATA3^-^/FOXA1^-^ (**Fig. 9E**), and CK5/6^-^/CK14^-^/CK19^-^ **(Fig. S9D)**. Unlike luminal breast epithelial cell derived tumors obtained after transformation with the same set of oncogenes ^22^, which metastasized to lungs, these tumors did not show extensive lung metastasis (**Fig. 9F and G**). Histologically, these tumors are metaplastic carcinomas, which comprise 0.08-0.2% of all breast neoplasms (**Fig. 9G**) ^19^.

### Normal breast and/or breast tumors of AA women have higher levels of phospho-STAT3

Since PZP-epithelial cell interaction resulted in elevated IL-6 expression *in vitro* and PZP cells are present at higher levels in the normal breast of AA women, it is expected that signals downstream of IL-6 should be higher in the breast tissues of AA women compared to EA women. Phosphorylation of STAT3 at residue S727 is one the major downstream signal upon binding of IL-6 to its receptor and S727 phosphorylated STAT3 promotes mesenchymal to epithelial transition of early disseminated cancer cells ^53^. We used phospho-STAT3 as a surrogate marker to determine IL-6 activity in the normal breast tissues and breast tumors of AA and EA women. Although cells in the KTB-normal contained lower levels of phospho-STAT3 as we could estimate on positivity, positivity was still higher with healthy breast tissues of AA women compared to EA women (**Fig. 10A-C**). A modestly higher levels of phospho-STAT3 were also observed in tumors of AA compared to EA women (**Fig. 10C**). Consistent with elevated levels of PZP cells in NATs compared to KTB-normal in EA women, phospho-STAT3 levels were higher in NATs compared to KTB-normal of EA women (**Fig. 10D**). Thus, genetic ancestry dependent variability in levels of stromal PZP cells has an impact on normal and cancerous breast biology.

**Figure 10:**
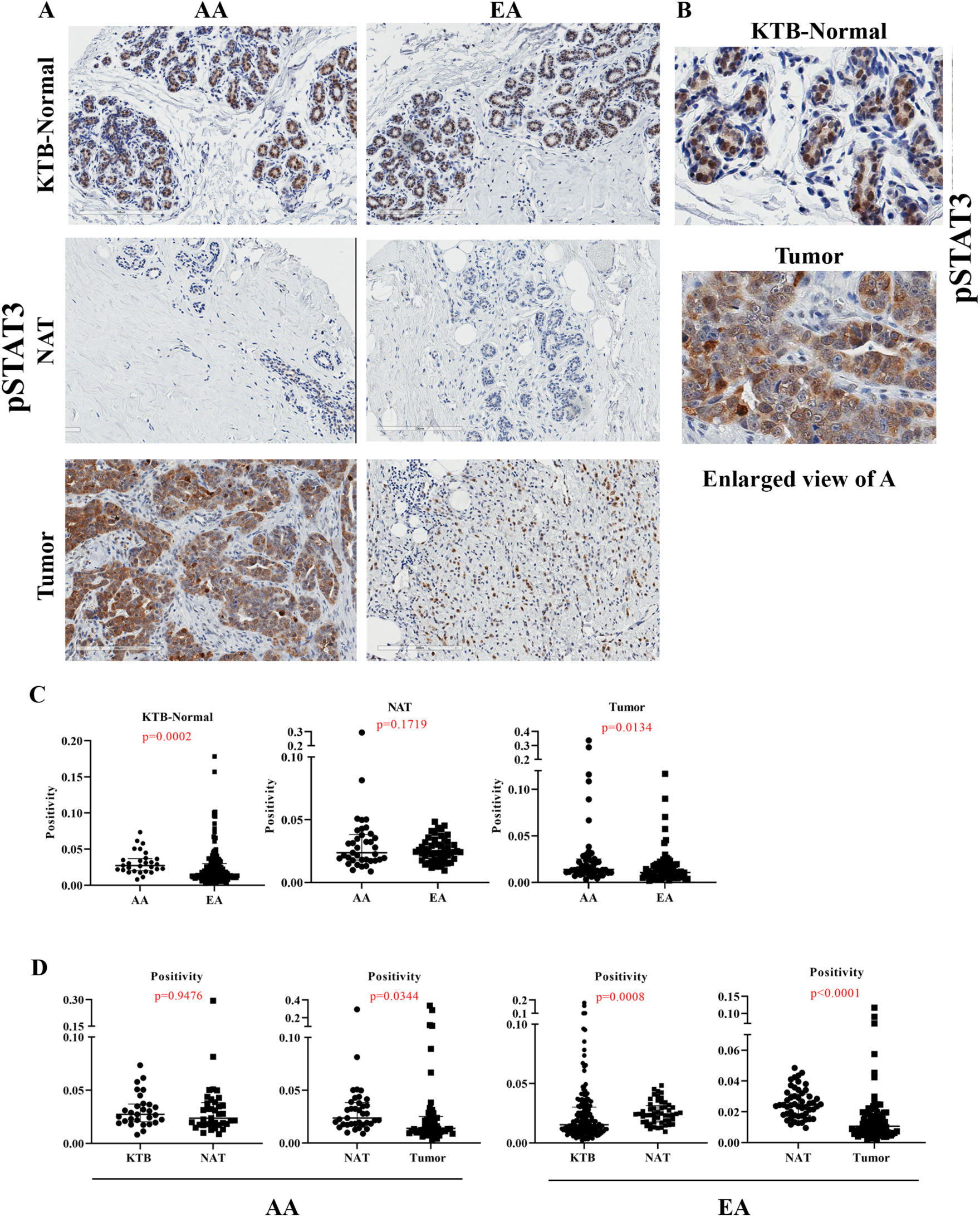
Phospho-STAT3 levels in KTB-normal, NATs, and breast tumors. (A) Representative IHC of pSTAT3 in KTB-normal, NATs, and/or tumors of women of AA, and EA. (B) Enlarged view of pSTAT3 expression in KTB-normal and tumor. (C) Differences in pSTAT3 positivity between KTB-normal, NATs and tumors of AA and EA women (D) Differences in pSTAT3 positivity between KTB-normal and NATs and between NATs and tumors of AA and EA women.

## DISCUSSION

At the most recent American Cancer Society estimate, breast cancer mortality rate in AA women is significantly higher compared to EA women, while breast cancer incidence in AA women is lower ^54^. This difference cannot be explained solely by differences in socioeconomic status or access to health care. This suggests that the biology of normal breast epithelial cells between these two groups differs, which may contribute to altered susceptibility to tumor initiation, progression and/or metastasis. This study focused on one particular stromal cell type, which can contribute to genetic ancestry dependent variability in breast biology.

Cell surface marker profiling and transdifferentiation studies indicate similarities between PZP cells and tissue resident mesenchymal stem cells, which have transdifferentiating and regenerative capacity ^55^. PZP cells contain three subpopulations, CD90^‒^/CD73^+^, CD90^low^/CD73^+^ and CD90^high^/CD73^+^. These cells also differentiate into adipocyte and osteogenic lineages. Similar cell type in the mouse mammary gland functions as adipogenic progenitor and differentiates into luminal and basal epithelia during mammary morphogenesis ^15^. We observed epithelial differentiation of PZP cells only upon transformation by HRas^G12V^. In this respect, PZP cells are the breast counterpart of PDGFRα^+^ intestinal stromal progenitors or fibroadipogenic progenitors (FAPs) of the skeletal muscle. FAPs cells are multipotent, and are capable of adipogenesis, fibrogenesis, osteogenesis and chondrogenesis ^38, 55^. While intestine and skeletal muscle undergo extensive regeneration upon injury, normal breast undergoes regeneration or modification during the pregnancy-lactation-involution cycle. Therefore, it is not surprising that such stromal cells exist in the breast.

PZP cells also displayed stromal fibroblast-like properties. Morsing M. et al. reported that a distinct fibroblast lineage in the human breast, which supports luminal epithelial progenitor growth and possessing adipogenic capacity, gathers around terminal ductal lobular units (TDLUs) ^28^. These cells are likely derived from lobular origin, which is evident by the enrichment of CD105^high^/CD26^low^ population. We found that PZP cells stably maintain the CD105^high^/CD26^low^ phenotype with extended culture. Lobular-like fibroblasts regulate epithelial morphogenesis and differentiation typical of the TDLU through TGFβ signaling pathway ^27, 28^. Early myoepithelial progenitors are susceptible to cues from lobular-like and interlobular-like fibroblasts in terms of luminal differentiation repertoire ^27, 28^. Since PZP cells expressed several secretory factors, more than luminal cells, few of these factors could provide specific cues to luminal cells.

Co-culture experiments provided a clue to how alterations in number of PZP cells in the breast could influence the breast microenvironment. For example, we observed a higher level of TAGLN expression under co-culture conditions compared to either cell types cultured separately. Recent studies have shown that TAGLN localizes in the cytoplasm of benign and malignant tumor cells in the breast tissue ^36^. It is strongly expressed in the cytoplasm of myoepithelial and fibroblastic cells of the benign breast tissue; however normal luminal cells are predominately negative or have a very weak cytoplasmic expression. TAGLN differentially expressed among molecular breast cancer subtypes and predominantly upregulated in higher grade triple negative breast cancers ^36^. Expression of TAGLN is positively correlated with high Ki-67 and low ER and PR status. TAGLN expression is found in 33% of the invasive carcinoma ^36^. Other studies showed that TAGLN is directly associated with migratory and invasive ability of tumorigenic cells and TAGLN overexpression increases the invasiveness of both tumorigenic and nontumorigenic cells ^56^.

PZP cell-epithelial cell interaction leading to elevated local production of IL-6 could have a profound effect on the tissue microenvironment. In this respect, genetic ancestry-dependent variability in breast tumor immune microenvironment has already been demonstrated, particularly in relation to CD8+ T-cell exhaustion ^57^. IL-6 is pleiotropic cytokine that contributes to EMT and drives metastasis in ER^+^ breast cancer ^58^. The IL-6/JAK/STAT3 pathway plays a major role in tumor growth and development of several human cancers. Previous studies report a higher levels of IL-6 in chronic inflammatory conditions, such as inflammatory bowel disease, rheumatoid arthritis, and in various cancers including breast cancer ^59^. IL-6 activates the JAK/STAT3 signaling, which drives the tumor proliferation, survival, invasiveness, angiogenesis and immunosuppression ^60^. Consistent with the possibility of genetic ancestry-dependent variability in PZP cells causing elevated local IL-6 levels and downstream signaling, we observed elevated phospho-STAT3 levels in normal and tumors of AA women compared to EA women.

Co-culturing of epithelial cells with PZP cells caused changes in the phenotype of epithelial cells with a fraction of luminal progenitor cells acquiring basal like properties. Recent studies have suggested that luminal cells with basal cell properties are enriched for gene signatures of basal-like breast cancers and number of cells with this property increase with aging ^17^. Whether locally produced IL-6 is responsible for luminal cells acquiring basal phenotype remains to be determined although there is evidence in the literature for IL-6 conferring plasticity to epithelial cells ^61^. Currently, at least four IL-6 targeting drugs are either approved or in clinical trial ^62^. It remains to be determined whether IL-6 targeting drugs are more effective against breast tumors of AA women compared to EA women.

Metaplastic carcinoma of the breast, although rare, is more common in AA women compared to EA women ^13^. Our results indicate that PZP cells are the cell-of-origin of this cancer type. Identifying cell-of-origin allows further evaluation of signaling pathways activated in this cancer type and eventually development of targeted therapies. Metaplastic carcinomas of the breast are CD49f^+^/EpCAM^-^ ^63^ and can be subdivided into metaplastic, claudin-(META-CLOW) and metaplastic-squamous, and metaplastic-non-CLOW, non-Squamous. PZP cells acquired CD49f^+^/EpCAM^-^ phenotype upon transformation and tumors derived from these cells displayed squamous features, indicating PZP cells are the cell-of-origin of metaplastic-squamous subtype.

In summary, this study identified and established PZP cell lines from healthy breasts of AA women and these cells phenotypically resemble tissue resident mesenchymal stromal/stem cells with transdifferentiation capabilities and can regulate signaling in neighboring epithelial cells. Although we identified only two factors, TAGLN and IL-6, whose levels in the tissue microenvironment could be influenced due to PZP-epithelial cell interaction, there may be additional signaling pathway activation as a consequence of this interaction and potentially influence the microenvironment. Because of the ability of PZP cells to influence signaling in epithelial cells and differences in their number based on genetic ancestry, signaling pathway in epithelial cells of the breast may show genetic ancestry-dependent variability in signaling under both normal and cancerous condition and drugs that target crosstalk between these two cell types can be new therapeutic agents. These drugs may show genetic ancestry-dependent variability in their potency. Therefore, in addition to including patients of different genetic ancestry in clinical trials, evaluation of outcomes of the clinical trial needs to take genetic ancestry into consideration. Some therapeutics, either as monotherapy or in combination, may be effective only in patients of specific genetic ancestry. Mechanistically, how PZP cell numbers are elevated in healthy breasts of AA women is unknown. FAPs, which are similar to PZP cells, increase in numbers in skeletal muscle of individuals with type II diabetes ^25^. Whether type II diabetes, which is more common in AA than EA ^64^, is responsible for increased PZPs in the breasts of AA women remains to be determined.

## ACKNOWLEDGMENTS

We thank members of the Indiana University Simon Comprehensive Cancer Center (IUSCCC) flow cytometry core, animal facility, and tissue procurement cores and the Susan G Komen Tissue Bank for various tissues and reagents. We also thank members of the Komen Tissue Bank and IUSCCC tissue bank including Jill Henry, Mary Cox, Rana German, Rockey Pam, and Julia von Arx for their work in generating TMA and associated data. We thank Dr. Natascia Marino for contributing part of EA TMA. We thank countless number of women for donating their breast tissue for research purpose as well as volunteers who facilitated tissue collection. This work is supported by DOD-W81XWH-15-1-0707, DOD-W81XWH-20-1-0577 and Susan G. Komen for the Cure (DRS20645418) to HN. Susan G. Komen for the Cure, Breast Cancer Research Foundation and Vera Bradley Foundation for Breast Cancer Research provide funding support to Komen Normal Tissue Bank.

## AUTHOR CONTRIBUTIONS

Conception and design: BK, HN Development of methodology: BK, PBN, LAB, CJT, AMS, HN Acquisition of data: BK, KB, PBN, MMG, RJA, MS, HN Analysis and interpretation of data: BK, KB, PBN, GS, SKA, HN Writing, review and/or revision of the manuscript: BK, HN Administrative, technical, or material support: HN, AMS, PBN Study supervision: HN

## DATA AVAILABILITY

This manuscript does not report any high throughput data. All reagents created as part of the manuscript will be distributed upon meeting requirements of the Indiana University.

