## Supplementary figures and a table for "Influence of genetic ancestry on breast stromal cells provides biologic basis for increased incidence of metaplastic breast cancer in women of African descent"

**Supplementary data.**

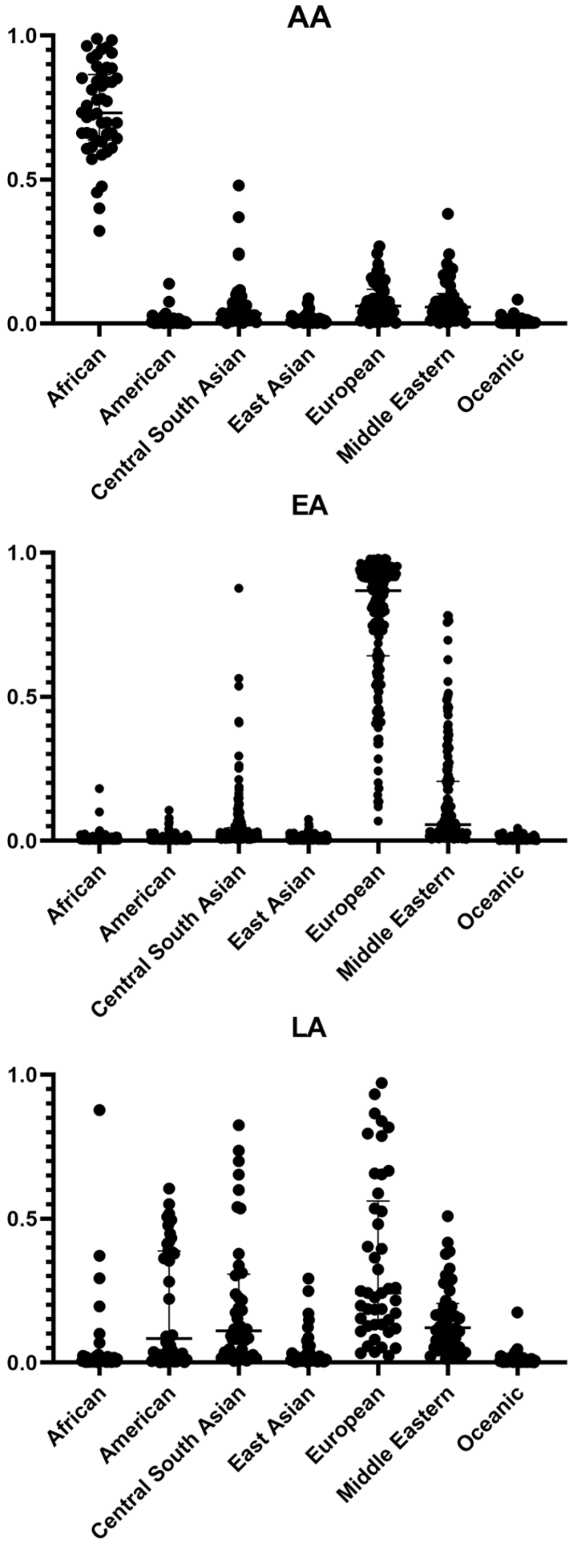
AA
EA
LA
1.0
0.5
0.0
African
American
Central South Asian
East Asian
European
Middle Eastern
Oceanic

Figure S1

**Figure S1: Genetic ancestry marker distribution patterns of donors whose breast tissues were used to generate TMA.**

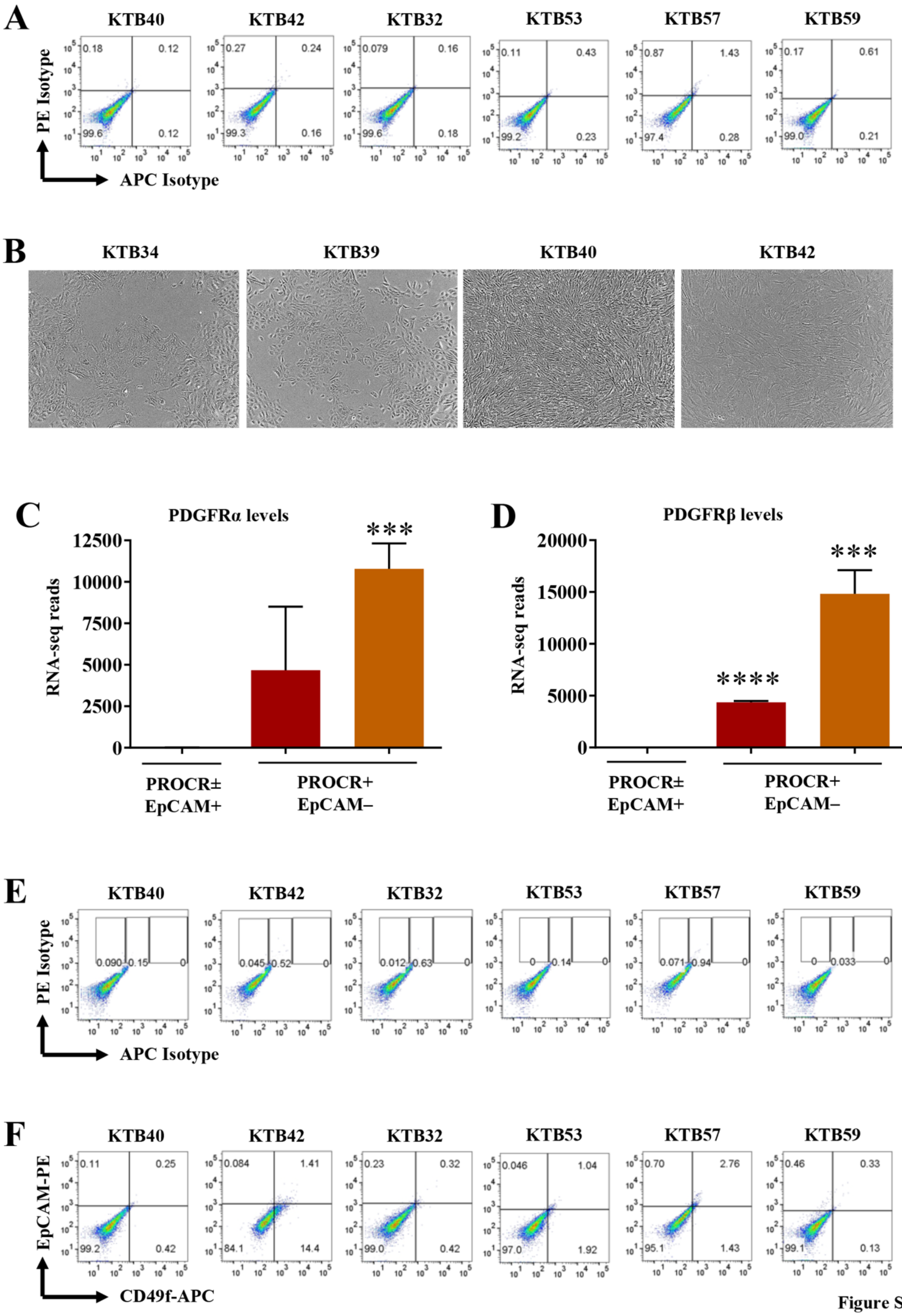
A
KTB40
KTB42
KTB32
KTB53
KTB57
KTB59
PE Isotype
APC Isotype
B
KTB34
KTB39
KTB40
KTB42
C
PDGFRα levels
RNA-seq reads
***
PROCR±
EpCAM+
PROCR+
EpCAM–
D
PDGFRβ levels
RNA-seq reads
****
***
PROCR±
EpCAM+
PROCR+
EpCAM–
E
KTB40
KTB42
KTB32
KTB53
KTB57
KTB59
PE Isotype
APC Isotype
F
KTB40
KTB42
KTB32
KTB53
KTB57
KTB59
EpCAM-PE
CD49f-APC

Figure S2

**Figure S2: PROCR$^{+}$/ZEB1$^{+}$ cells express PDGFRα**. (A) APC and PE isotypes controls of cell lines characterized by flow cytometry. These staining patterns were used to draw quadrants. (B) Phase contrast images representing the epithelial morphology of human telomerase gene (hTERT) immortalized breast epithelial cells (KTB34 and KTB39) and fibroblast-like features of KTB40 and KTB42. (C) PDGFRα mRNA level in PROCR$^{+}$/EpCAM$^{-}$ cells compared to PROCR$^{\pm}$/EpCAM$^{+}$ cells. (D) PDGFRβ mRNA level in PROCR$^{+}$/EpCAM$^{-}$ cells compared to PROCR$^{\pm}$/EpCAM$^{+}$ cells. (E) APC and PE isotypes controls of cell lines characterized by flow cytometry. These staining patterns were used to draw quadrants. (F) PZP Cell lines were stained with CD49f and EpCAM antibodies to demonstrate lack of breast stem, luminal progenitor, and mature/differentiated cell features. ***$p<0.001$, ****$p<0.0001$ by ANOVA.

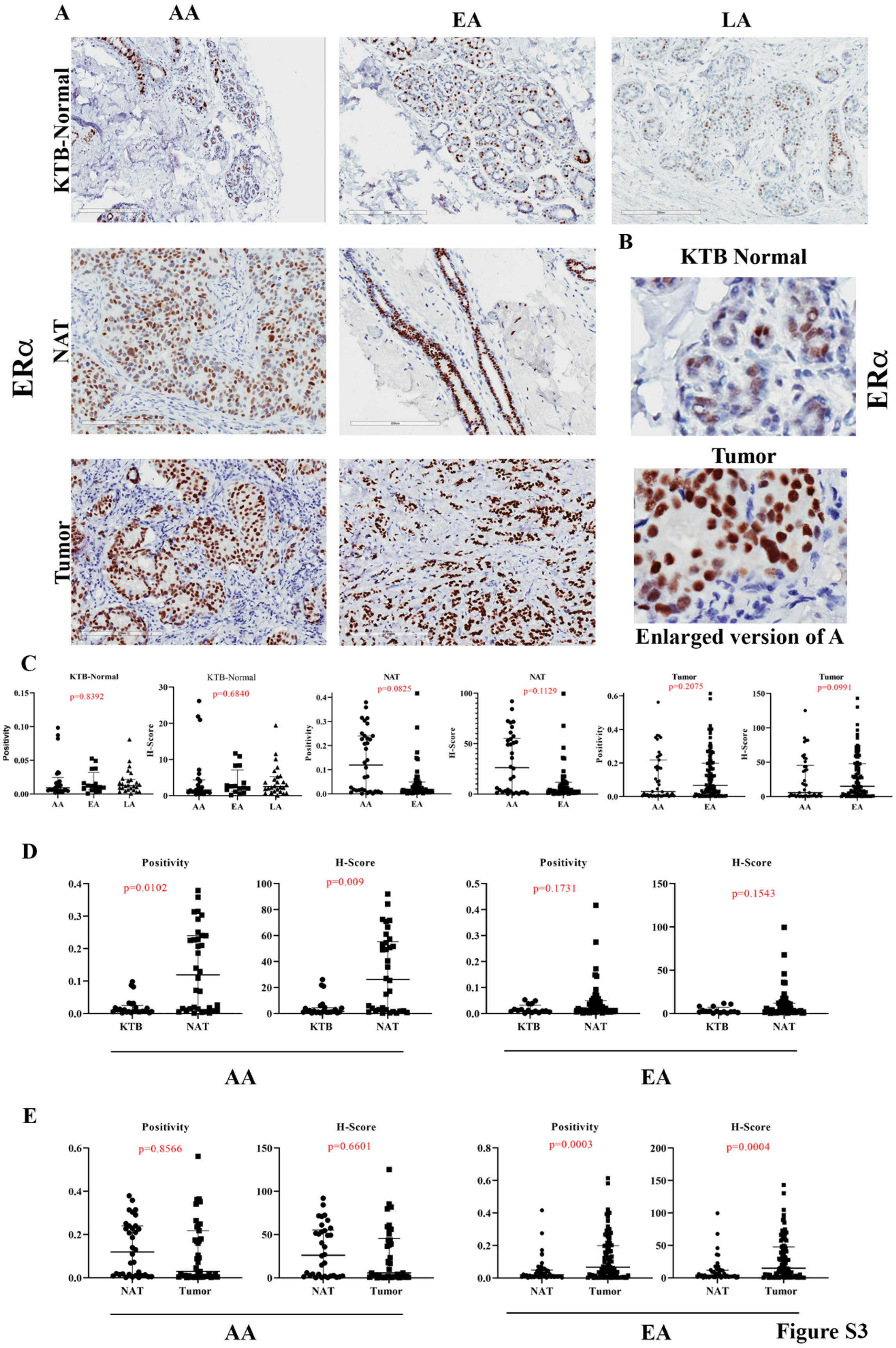
A
AA
EA
LA
KTB-Normal
NAT
Tumor
ERα
B
KTB Normal
Tumor
ERα
Enlarged version of A
C
KTB-Normal
p=0.8392
KTB-Normal
p=0.6840
NAT
p=0.0825
NAT
p=0.1129
Tumor
p=0.2075
Tumor
p=0.0991
Positivity
H-Score
D
Positivity
p=0.0102
H-Score
p=0.009
Positivity
p=0.1731
H-Score
p=0.1543
KTB
NAT
AA
EA
E
Positivity
p=0.8566
H-Score
p=0.6601
Positivity
p=0.0003
H-Score
p=0.0004
NAT
Tumor
AA
EA

Figure S3

**Figure S3: ERα expression pattern in KTB-normal, NATs, and breast tumors**. (A) Representative IHC of ERα in KTB-normal, NATs, and tumors of women of AA, EA, and LA women. (B) Enlarged view of ERα expression in KTB-normal and tumor. (C) Differences in ERα expression (positivity and H-score) between KTB-normal of AA, EA, and LA women. Differences in ERα expression (positivity and H-score) between KTB-normal, NATs and/or tumors of AA, EA, and LA women. (D) Differences in ERα expression (positivity and H-score) between KTB-normal and NATs in AA and EA women. (E) Differences in ERα expression (positivity and H-score) between NATs and tumors in AA and EA women.

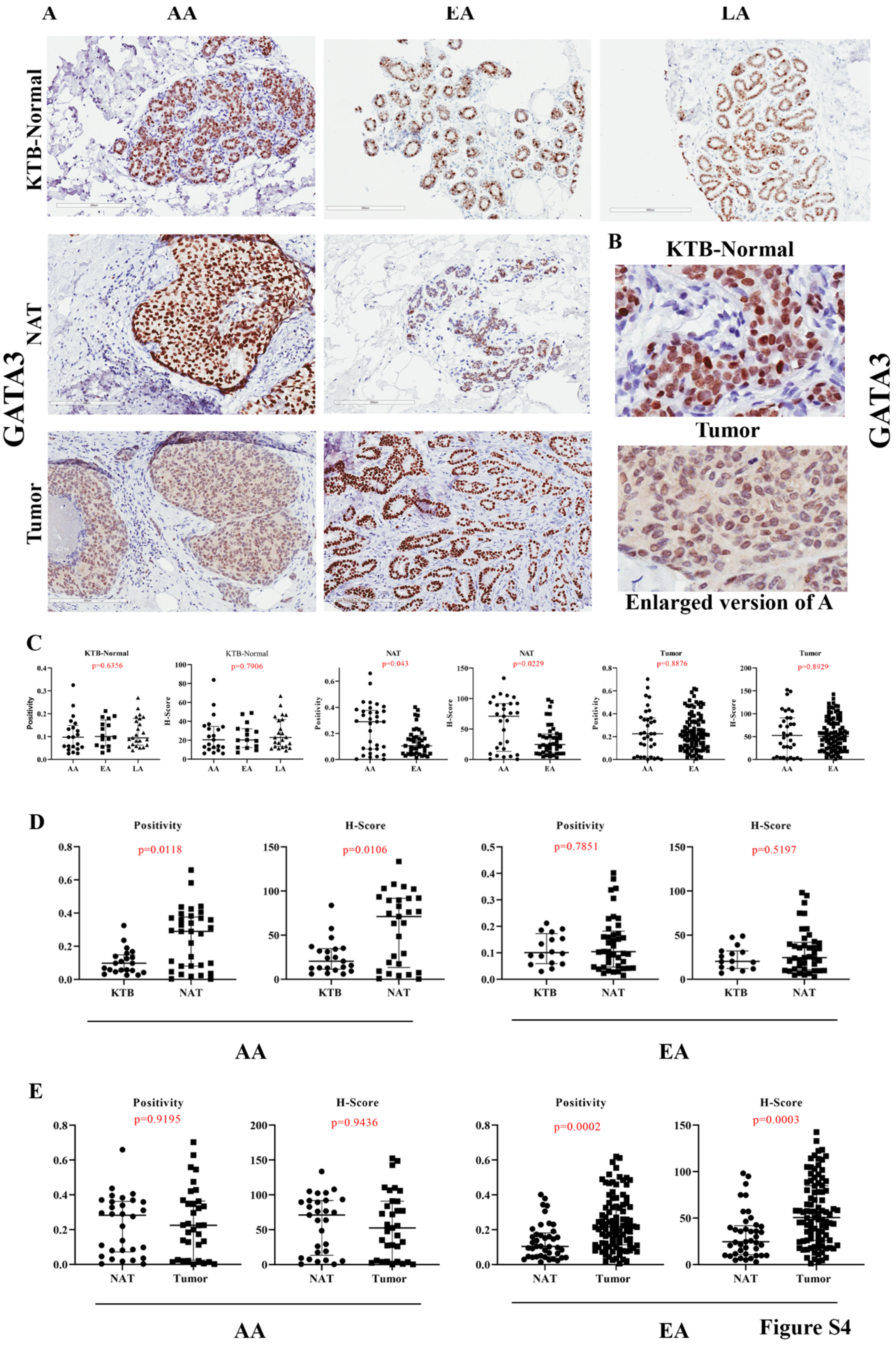
A
AA
EA
LA
KTB-Normal
NAT
Tumor
GATA3
B
KTB-Normal
Tumor
GATA3
Enlarged version of A
C
KTB-Normal
p=0.6356
KTB-Normal
p=0.7906
NAT
p=0.043
NAT
p=0.0229
Tumor
p=0.8876
Tumor
p=0.8929
Positivity
H-Score
AA
EA
LA
D
Positivity
p=0.0118
H-Score
p=0.0106
Positivity
p=0.7851
H-Score
p=0.5197
KTB
NAT
AA
EA
E
Positivity
p=0.9195
H-Score
p=0.9436
Positivity
p=0.0002
H-Score
p=0.0003
NAT
Tumor
AA
EA

Figure S4

**Figure S4: GATA3 expression pattern in KTB-normal, NATs, and breast tumors**. (A) Representative IHC of GATA3 in KTB-normal, NATs, and tumors of women of AA, EA, and LA women. (B) Enlarged view of GATA3 expression in KTB-normal and tumor. (C) Differences in GATA3 expression (positivity and H-score) between KTB-normal of AA, EA, and LA women. Differences in GATA3 expression (positivity and H-score) between KTB-normal, NATs and/or tumors of AA, EA, and LA women. (D) Differences in GATA3 expression (positivity and H-score) between KTB-normal and NATs in AA and EA women. (E) Differences in GATA3 expression (positivity and H-score) between NATs and tumors in AA and EA women.

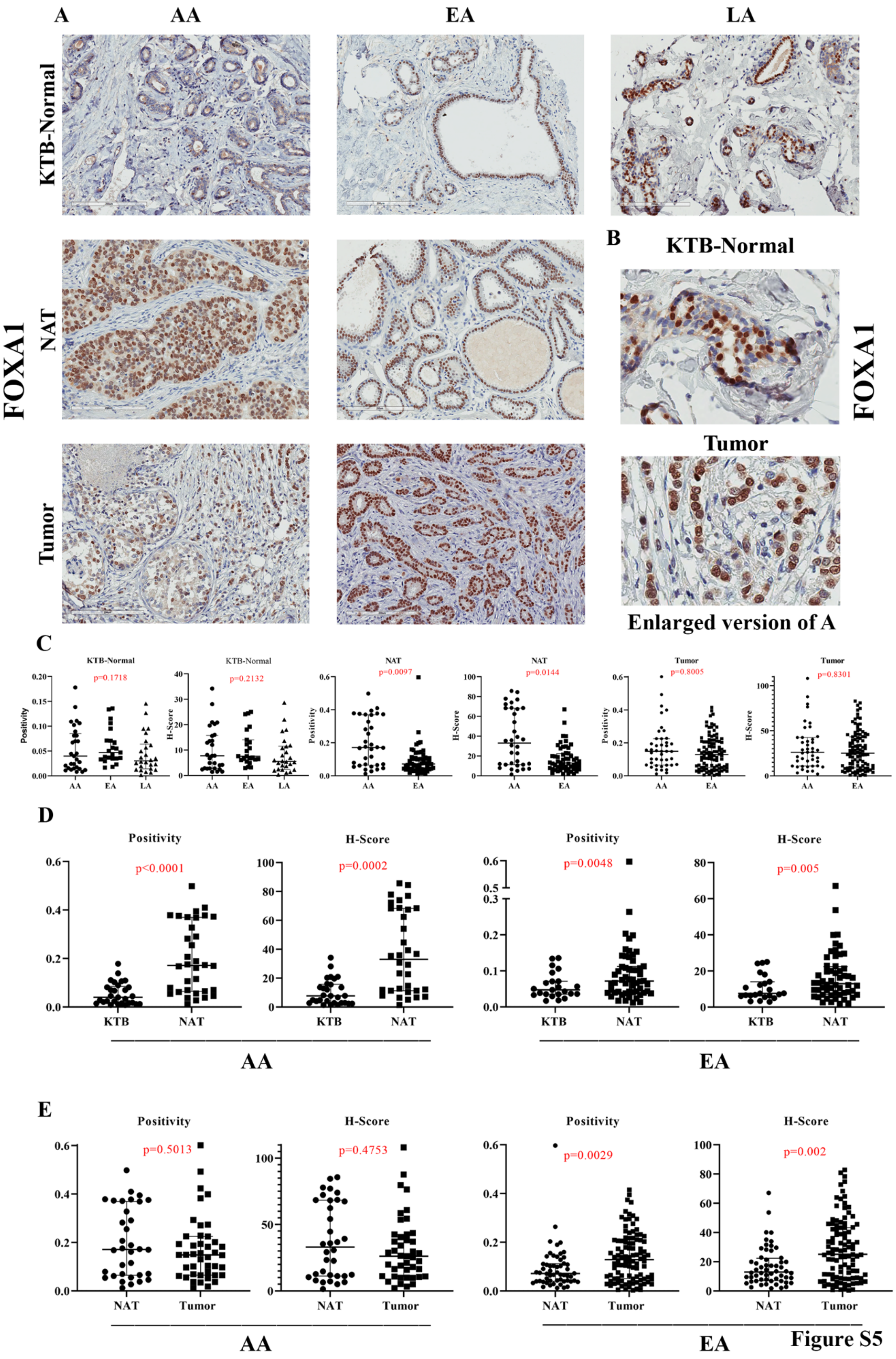
A
AA
EA
LA
KTB-Normal
NAT
Tumor
FOXA1
B
KTB-Normal
Tumor
FOXA1
Enlarged version of A
C
KTB-Normal p=0.1718
KTB-Normal p=0.2132
NAT p=0.0097
NAT p=0.0144
Tumor p=0.8005
Tumor p=0.8301
Positivity
H-Score
AA EA LA
AA EA
D
Positivity p<0.0001
H-Score p=0.0002
Positivity p=0.0048
H-Score p=0.005
KTB NAT
AA
EA
E
Positivity p=0.5013
H-Score p=0.4753
Positivity p=0.0029
H-Score p=0.002
NAT Tumor
AA
EA

Figure S5

**Figure S5: FOXA1 expression pattern in KTB-normal, NATs, and breast tumors**. (A) Representative IHC of FOXA1 in KTB-normal, NATs, and tumors of women of AA, EA, and LA women. (B) Enlarged view of FOXA1 expression in KTB-normal and tumor. (C) Differences in FOXA1 expression (positivity and H-score) between KTB-normal of AA, EA, and LA women. Differences in FOXA1 expression (positivity and H-score) between KTB-normal, NATs and/or tumors of AA, EA, and LA women. (D) Differences in FOXA1 expression (positivity and H-score) between KTB-normal and NATs in AA and EA women. (E) Differences in FOXA1 expression (positivity and H-score) between NATs and tumors in AA and EA women.

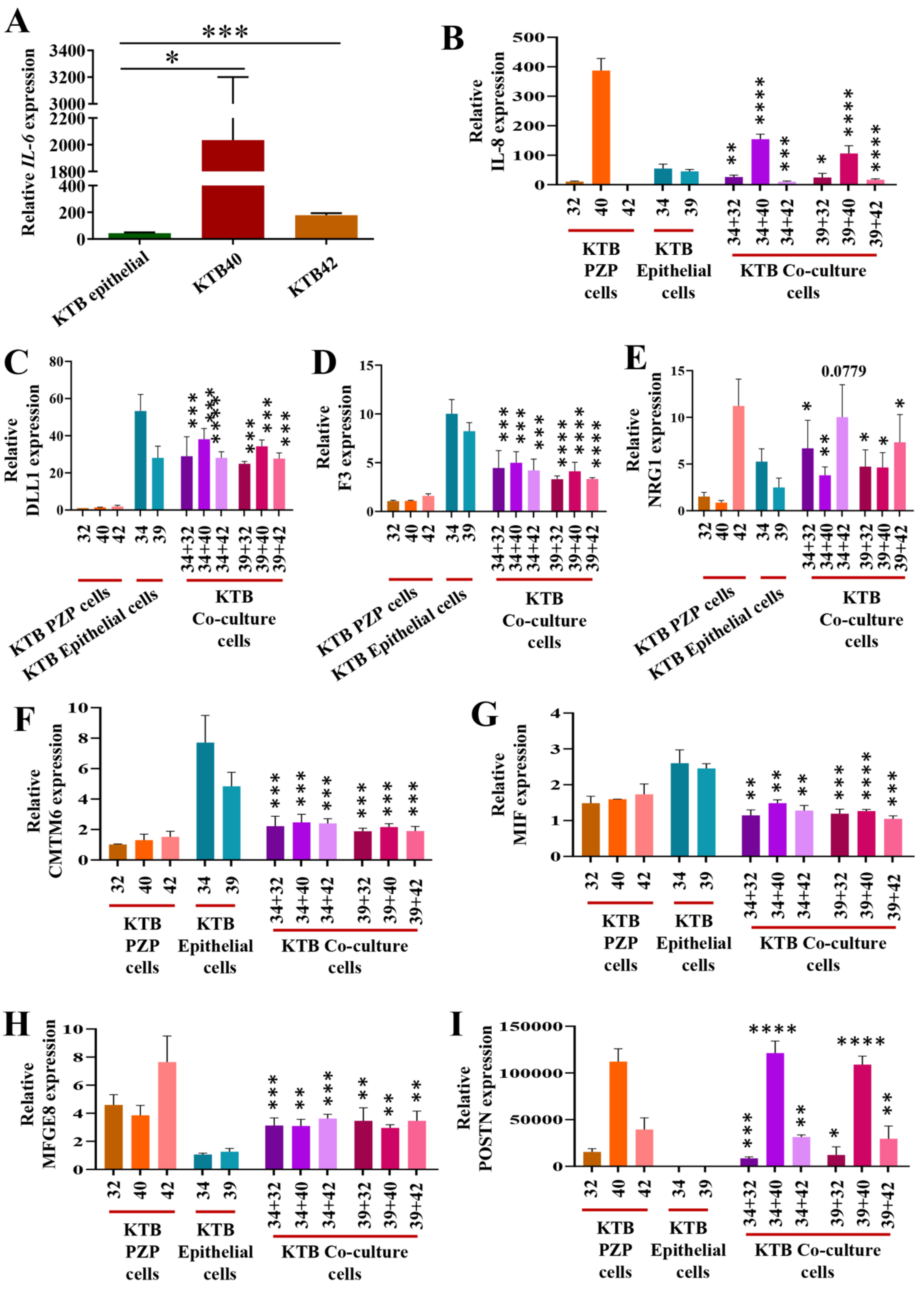
A
Relative IL-6 expression
KTB epithelial
KTB40
KTB42
B
Relative IL-8 expression
C
Relative DLL1 expression
D
Relative F3 expression
E
Relative NRG1 expression
0.0779
F
Relative CMTM6 expression
G
Relative MIF expression
H
Relative MFGE8 expression
I
Relative POSTN expression
KTB PZP cells
KTB Epithelial cells
KTB Co-culture cells

Figure S6

**Figure S6: Effect of co-culture of PZP and luminal progenitor cell lines on expression of genes**. (A) Relative mRNA level of IL-6 in KTB40, KTB42 and KTB-epithelial cells. Expression of IL-8 (B), DLL1 (C), F3 (D), and NRG1 (E), CMTM6 (F), MIF (G), MFGE8 (H), and POSTN (I) in PZP cell lines (KTB32, KTB40, KTB42), luminal progenitor (epithelial cells; KTB34, KTB39), and co-culture of PZP and luminal progenitor cell lines. Statistical significance (*p* values) was determined by comparing KTB32/40/42 (PZP cells), KTB34/KTB39 (epithelial cells) and respective co-cultured PZP + epithelial cells as indicated in the figure. *$p<0.05$, **$p<0.01$, ***$p<0.001$, ****$p<0.0001$ by ANOVA.

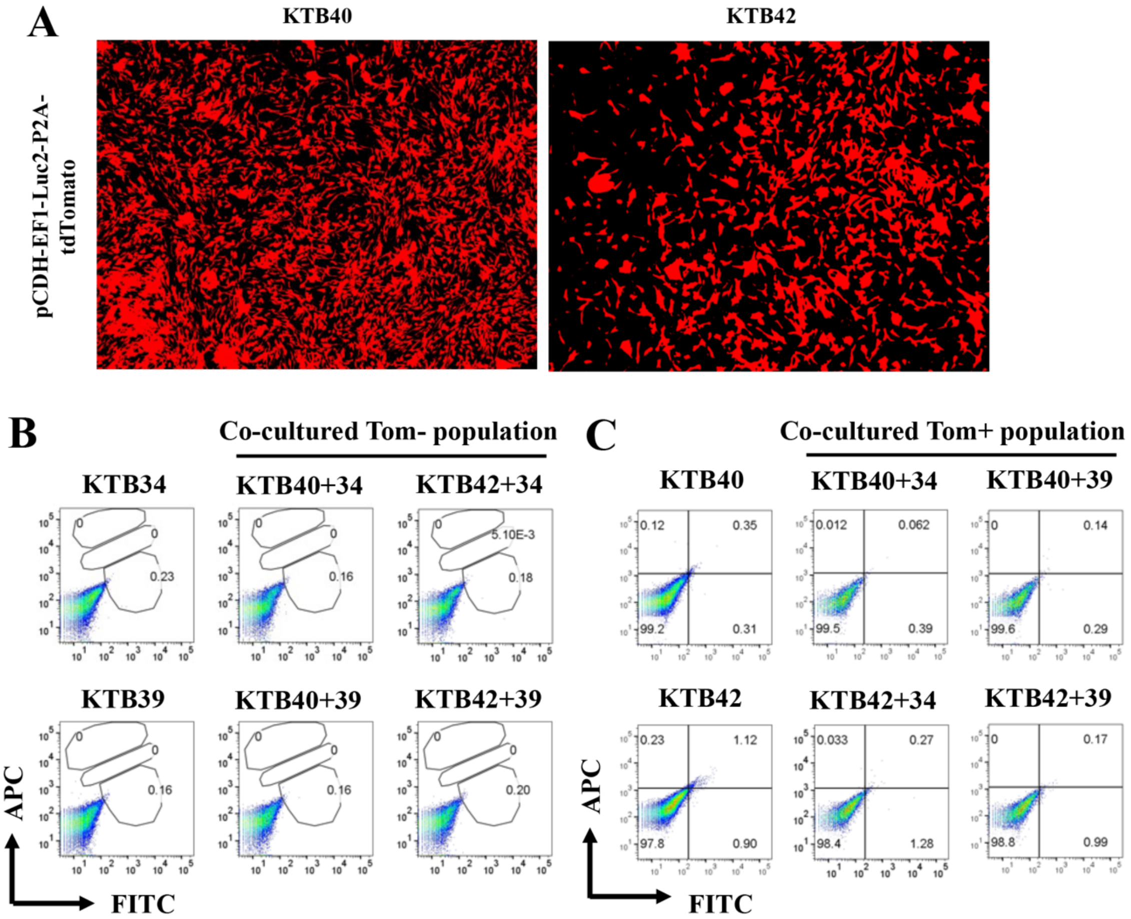
A
KTB40
KTB42
pCDH-EF1-Luc2-P2A-tdTomato
B
Co-cultured Tom- population
KTB34
KTB40+34
KTB42+34
KTB39
KTB40+39
KTB42+39
APC
FITC
C
Co-cultured Tom+ population
KTB40
KTB40+34
KTB40+39
KTB42
KTB42+34
KTB42+39
APC
FITC

Figure S7

**Figure S7: Generation of tomato red-labeled stable PZP cell lines**. (A) PZP KTB40 and KTB42 cells were labelled with tomato-red using pCDH-EF1-Luc2-P2A-tdTomato lentivirus and sorted the tomato-red positive cells by flow cytometry to generate the stable cell lines. (B and C) APC and FITC isotypes controls of cell lines characterized by flow cytometry. These staining patterns were used to draw quadrants.

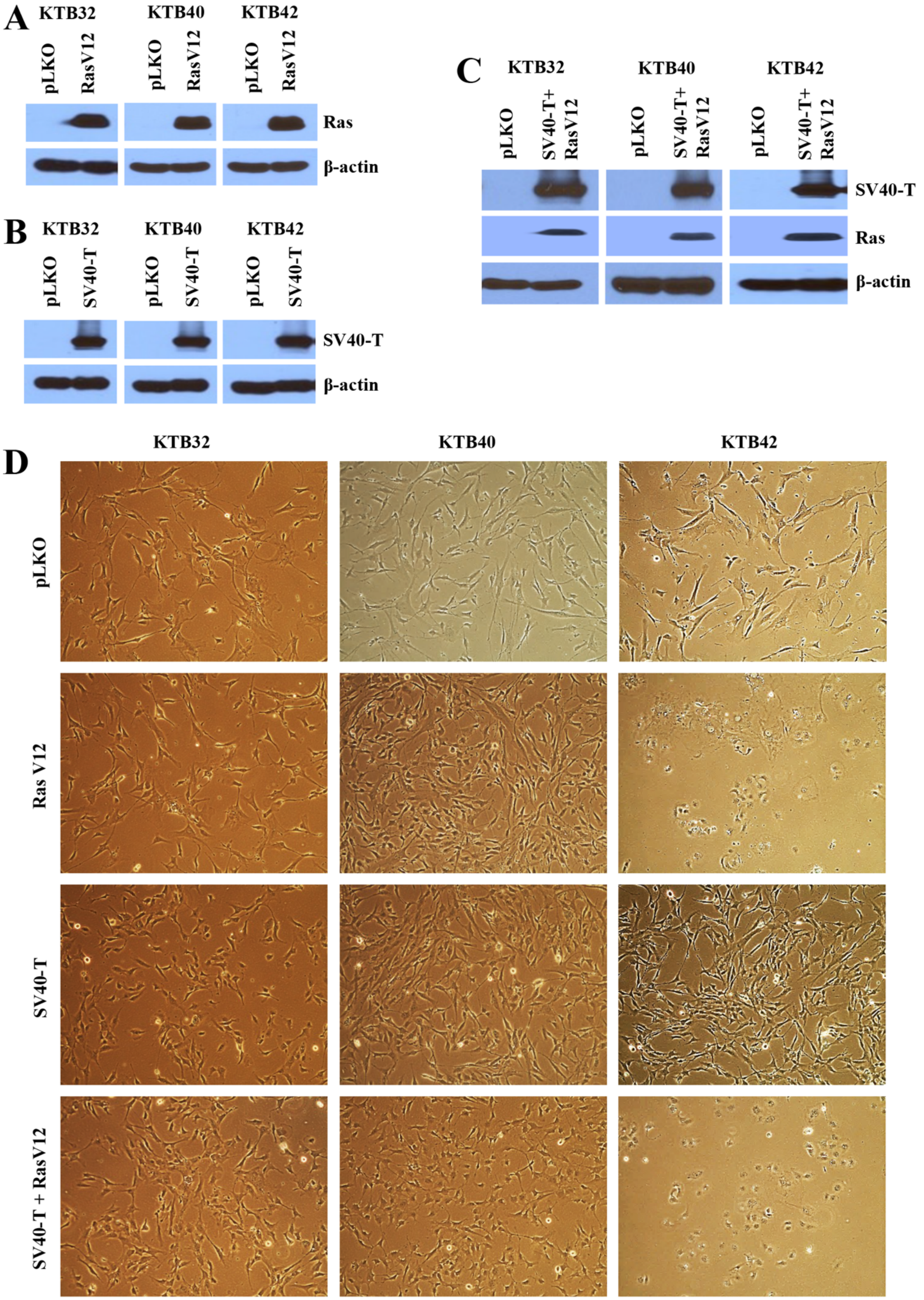
A
KTB32
KTB40
KTB42
pLKO
RasV12
Ras
β-actin
B
SV40-T
C
SV40-T+ RasV12
D
Ras V12
SV40-T + RasV12

Figure S8

**Figure S8: Generation of $HRas^{G12V}$, SV40-T/t antigen and $HRas^{G12V}$+ SV40-T/t antigen transformed PZP cell lines**. (A) Western blotting was used to detect overexpression of mutant Ras in KTB32, KTB40 and KTB42. (B) Western blotting was used to detect overexpression of SV40-T/t antigen in KTB32, KTB40 and KTB42. (C) Western blotting was used to detect overexpression of mutant Ras and SV40-T/t antigen in double transformed KTB32, KTB40 and KTB42. pLKO was used as a control cell line. β–actin was used as an internal control. (D) Phase contrast images showing morphology of KTB32, KTB40 and KTB42 transformed with $HRas^{G12V}$, SV40-T/t antigen and $HRas^{G12V}$+ SV40-T/t antigen.

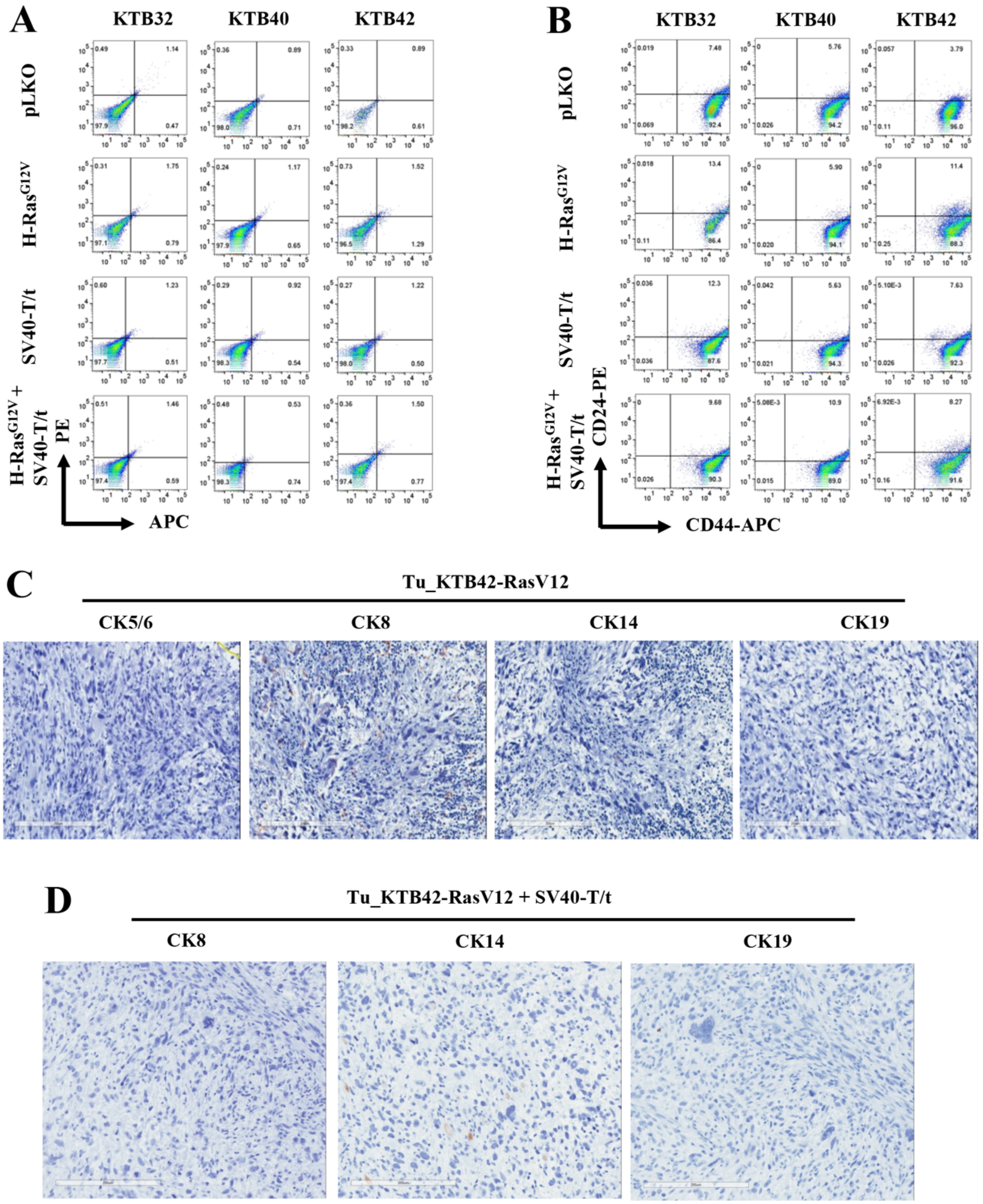
A
KTB32
KTB40
KTB42
pLKO
H-Ras^G12V
SV40-T/t
H-Ras^G12V +
SV40-T/t
PE
APC
B
KTB32
KTB40
KTB42
pLKO
H-Ras^G12V
SV40-T/t
H-Ras^G12V +
SV40-T/t
CD24-PE
CD44-APC
C
Tu_KTB42-RasV12
CK5/6
CK8
CK14
CK19
D
Tu_KTB42-RasV12 + SV40-T/t
CK8
CK14
CK19

Figure S9

**Figure S9: Characterization of transformed cell lines and tumors**. (A) APC and PE isotypes controls of cell lines characterized by flow cytometry. These staining patterns were used to draw quadrants. (B) CD44 and CD24 staining patterns of immortalized and transformed PZP (KTB32, KTB40 and KTB42) cell lines. (C) IHC analyses of cytokeratins CK5/6, CK14, CK8 and CK19 in tumors developed from the KTB42-HRas$^{G12V}$ transformed cells. Basal marker: CK5/6 and CK14, Luminal marker: CK8 and CK19. (D) IHC analyses of cytokeratins CK14, CK8 and CK19 in tumors developed from the KTB42-HRas$^{G12V}$+SV40-T/t antigen transformed cells.

**Table S1:** Histopathological features of tumors used in the TMA.

| | **AA** | **EA** |
|---|---|---|
| $ER^+$ | 29 | 104 |
| $PR^+$ | 23 | 88 |
| $Her2^+$ | 12 | 22 |
| $Node^+$ | 16 | 20 |
| Grade 1 | 1 | 10 |
| Grade 2 | 11 | 69 |
| Grade 3 | 23 | 36 |
| Stage 1 | 0 | 5 |
| Stage 2 | 9 | 17 |
| Stage 3 | 1 | 6 |
| Stage 4 | 1 | 1 |
| No data available | 5 | 1 |
| Total Samples | 49 | 136 |
